# TPClust: Temporal Profile-Guided Subtyping Using High-Dimensional Omics Data

**DOI:** 10.1101/2025.08.05.668514

**Authors:** Boyi Hu, Philip L. De Jager, David A. Bennett, Badri N. Vardarajan, Yuanjia Wang, Annie J. Lee

**Author notes:** Correspondence: Annie J. Lee.

## Abstract

Clustering is widely used to identify subtypes in heterogeneous populations, yet most approaches rarely integrate longitudinal phenotypic trajectories with high-dimensional molecular profiles, limiting their ability to resolve biologically and clinically meaningful heterogeneity in progressive diseases. We developed TPClust, a supervised, semi-parametric clustering method that integrates high-dimensional omics data with longitudinal phenotypes including outcomes and covariates for outcome-guided subtyping. TPClust jointly models latent subtype membership and longitudinal outcome trajectories using multinomial logistic regression informed by molecular features selected via structured regularization, along with spline-based regression to capture subtype-specific, time-varying covariate effects. Simulations demonstrate valid inference for time-varying effects and robust feature selection. Applied to transcriptomic profiles and longitudinal cognitive data from 1,020 older adults in the Religious Orders Study and the Rush Memory and Aging Project, TPClust identified four aging subtypes including intermediate subtypes not captured by unimodal approaches with distinct cognitive trajectories, time-varying risk profiles, clinical and neuropathological features, and multimodal molecular signatures.

## Introduction

Clustering is widely used to identify subgroups within heterogeneous populations, uncovering molecular or phenotypic subtypes that can inform disease mechanisms and precision medicine. The advent of high-throughput technologies has enabled large-scale profiling of molecular data, such as transcriptomics and proteomics. When integrated with longitudinal phenotypes such as cognitive or biomarker trajectories, these profiles support systematic investigation of temporal dynamics in disease progression. However, most clustering methods do not jointly model high-dimensional molecular features and longitudinal phenotypes, limiting their ability to identify biologically and clinically meaningful heterogeneity in progressive conditions.

Traditional unsupervised clustering methods, such as K-means clustering (Forgy, 1965; Sahelijo et al., 2024; Lee et al., 2023), are typically applied to a single data modality—most often molecular data—and do not jointly model molecular and phenotypic information. While useful for detecting molecular differences, the resulting subtypes often lack clinical interpretability and may reflect technical noise or disease-irrelevant structure. In Alzheimer’s disease (AD), where clinical manifestations and risk factor effects (e.g., *APOEε*4, early-onset diabetes) evolve with age (Tang et al., 1998; Bellou et al., 2020; Amidei et al., 2021), unsupervised clustering has been applied to transcriptomic profiles or co-expression networks (Milind et al., 2020; Neff et al., 2021; Lee et al., 2023; Sahelijo et al., 2024). However, without incorporating phenotypic progression, these subtypes may miss clinically relevant patterns, highlighting the need for integrative methods that jointly model high-dimensional omics, longitudinal phenotypic data, and time-varying covariate effects.

Supervised clustering methods address this gap by incorporating phenotypic outcomes into subtype discovery. Early approaches targeted binary or cross-sectional outcomes with low-dimensional molecular data (Guo et al.,2006), and later extensions accommodated high-dimensional molecular data using penalized regression (House et al., 2006; Meng et al., 2022), though still limited to static outcomes without covariate adjustment. Recent methods, such as ogClust (Li et al., 2024), allow covariate adjustment with high-dimensional omics data but remain restricted to cross-sectional outcomes. Longitudinal mixture models (Proust-Lima et al., 2014; Sun et al., 2019) capture longitudinal outcome trajectories but assume low-dimensional features and time-invariant covariate effects.

Here, we developed Temporal Profile-guided Clustering (TPClust), a supervised, semi-parametric clustering method that integrates longitudinal outcomes with high-dimensional molecular data to identify biologically and clinically meaningful subtypes. TPClust jointly models subtype membership using omics features and outcome trajectories with spline-based regression to capture subtype-specific, time-varying covariate effects on longitudinal cognitive outcomes (Fig. 1). Structured regularization facilitates robust feature selection in the high-dimensional omics features. Model parameters are estimated via an expectation-maximization algorithm, and inference for time-varying effects is conducted using multiplier bootstrap (Van Der Vaart and Wellner, 1996). Simulations demonstrate valid inference and robust feature selection performance. The TPClust workflow proceeds in three stages: (1) model fitting to estimate subtype memberships, informative omics features, and dynamic covariate effects; (2) interpretation of subtype-specific time-varying effects to reveal temporal and clinical heterogeneity; and (3) downstream association analyses with clinical traits and biomarkers to characterize biological distinctions across subtypes. Applied to transcriptomic and longitudinal cognitive data from the Religious Orders Study and Rush Memory and Aging Project (ROSMAP), TPClust identifies four aging subtypes with distinct cognitive trajectories, risk factor dynamics, neuropathology, and multimodal molecular signatures.

**Figure 1:**
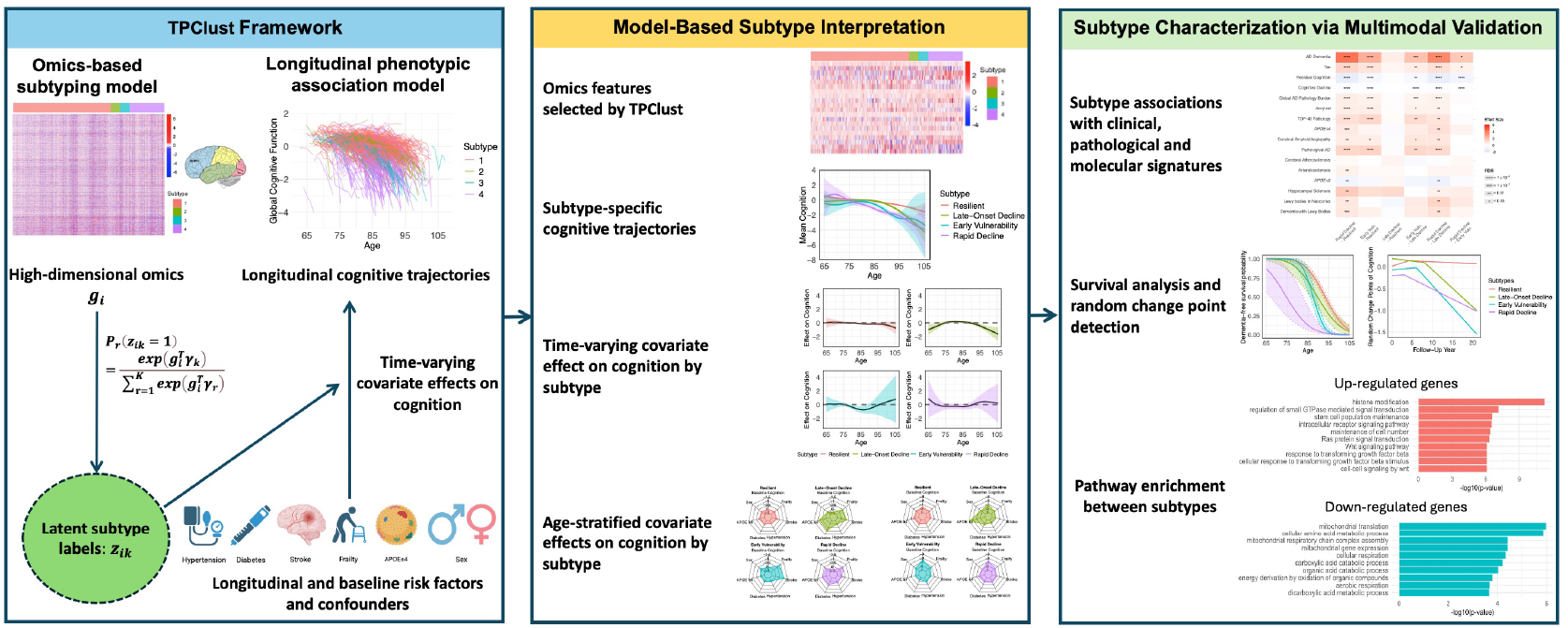
TPClust framework for outcome-guided subtyping. TPClust is a joint modeling method that integrates high-dimensional omics and longitudinal phenotypic data to identify biologically and clinically meaningful subtypes. The workflow comprises two components (left): an omics-based subtyping model that assigns latent subtype membership via regularized multinomial logistic regression, and a longitudinal association model that estimates subtype-specific, time-varying effects of covariates on cognitive function using B-spline regression. The model was applied to the ROSMAP cohort to identify TPClust-derived subtypes. Subtype interpretation (middle) includes transcriptomic profiles of subtype-discriminating features, estimated cognitive trajectories, time-varying covariate effects on cognition, and age-stratified summaries of covariate effects on cognition. Subtype characterization (right) integrates multimodal analyses, including associations with clinical, pathological, and molecular traits, survival analysis with change-point detection, and subtype-specific pathway enrichment.

## Results

### TPClust enables high-dimensional subtyping through time-varying covariate modeling

TPClust introduces a unified probabilistic method for supervised subtyping that simultaneously models (1) high-dimensional molecular features that contribute to subtypes and (2) longitudinal phenotypes, including smooth outcome trajectories and subtype-specific, time-varying covariate associations (Fig.1). This method addresses limitations of existing subtyping methods that assume time-invariant effects, enabling the identification of clinically meaningful subtypes with distinct progression dynamics and evolving risk profiles.

The model comprises two core components. Subtype membership is represented by a latent categorical variable, with assignment probabilities modelled through multinomial logistic regression informed by omics features. Structured feature selection is performed using regularized estimation using a combination of LASSO (Tibshirani, 1996), group LASSO (Meier et al., 2008), and sparse group LASSO (Simon et al., 2013) penalties, allowing integration of genome-wide and pathway-informed information while accommodating complex correlation structures. Longitudinal outcomes are modelled using semi-parametric regression with subtype-specific, time-varying covariate effects represented by B-splines. This allows for flexible modelling of nonlinear trajectories and dynamic associations across time.

Model parameters are estimated using a modified expectation-maximization algorithm (Meng and Van Dyk, 1997) that iteratively updates latent subtype assignments and associated parameters. Each iteration alternates between computing posterior probabilities of subtype membership, given observed longitudinal data, omics features and current estimates, and re-estimating parameters based on these updated assignments. Regularized optimization is applied to the expected complete-data likelihood, enabling simultaneous selection of informative features and estimation of time-varying covariate effects.

For inference on time-varying covariate–outcome associations, TPClust employs a multiplier bootstrap procedure (Van Der Vaart and Wellner, 1996) that accounts for within-subject correlation in longitudinal data. Unlike standard bootstrap methods that resample entire subjects and may fail to capture intra-subject dependencies, the multiplier bootstrap perturbs individual likelihood contributions by applying random weights. Model parameters are re-estimated under each perturbed likelihood, and the resulting empirical distribution of the time-varying coefficients is used to construct pointwise confidence intervals using normal approximation.

### Simulation studies validate TPClust’s inference and feature selection performance

To evaluate the performance of TPClust in recovering latent subtypes and modeling dynamic covariate effects, we simulated longitudinal data under a supervised mixture regression method (see Methods). Each dataset consisted of three subtypes assigned using a multinomial logistic model, with subtype probabilities determined by a subset of high-dimensional omics features comprising both informative and noise variables. Conditional on subtype, longitudinal outcomes were generated using smooth B-spline functions to represent subtype-specific, time-varying covariate effects, mimicking heterogeneous progression trajectories.

We considered two simulation scenarios, each replicated 100 times, with sample sizes of *n* = 500 and *n* = 1, 000. In Scenario 1, we included 10 informative and 100 noise features; in Scenario 2, dimensionality was increased to 20 informative and 500 noise features. To evaluate inference accuracy for the time-varying covariate effects, we computed the empirical coverage of multiplier bootstrap–derived pointwise confidence intervals. Across both scenarios and sample sizes, coverage rates closely matched the nominal 95% level, supporting the validity of the inference procedure (Fig. 2a). Estimation error, measured using integrated mean squared error, declined with increasing sample size, indicating improved trajectory estimation accuracy (Fig. 2b).

**Figure 2:**
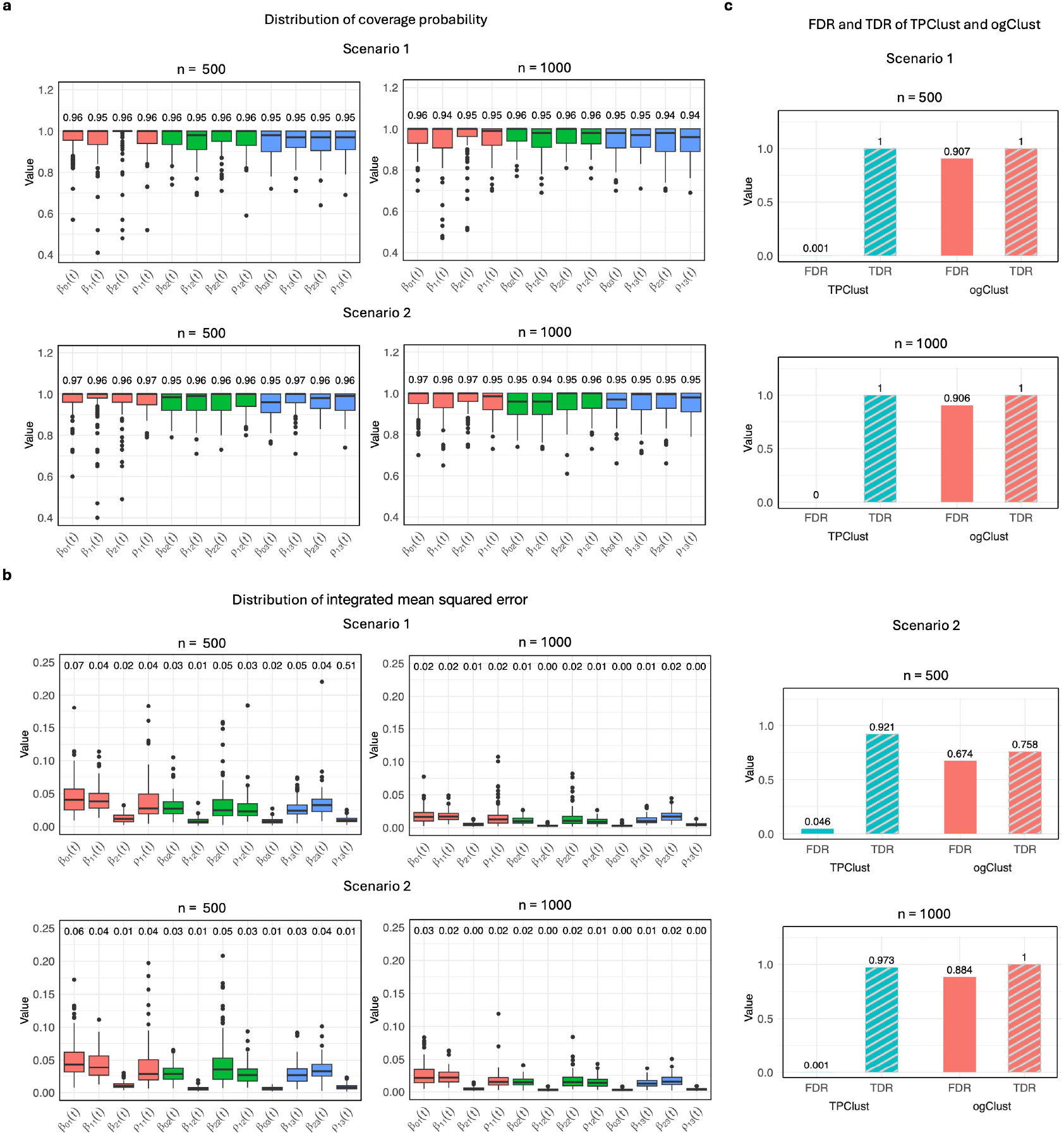
Simulation studies validate TPClust’s inference accuracy and feature selection performance. Two scenarios were evaluated: Scenario 1 (10 informative, 100 noise features) and Scenario 2 (20 informative, 500 noise features), each replicated 100 times with *n* = 500 and *n* = 1,000. a, Empirical coverage of 95% confidence intervals for time-varying effects using multiplier bootstrap, showing nominal coverage across all settings. b, Integrated mean squared error (IMSE) of estimated effects, demonstrating improved estimation accuracy with larger sample sizes and robustness across both scenarios. c, Feature selection results showing TPClust outperforms ogClustGM with lower FDR and higher TDR across all settings.

We next compared TPClust’s feature selection performance against ogClustGM (Li et al., 2024), a penalized Gaussian mixture model that supports high-dimensional inputs but assumes static covariate effects. In both scenarios, ogClustGM exhibited elevated false discovery rates (FDR), frequently selecting noise features. By contrast, TPClust consistently achieved higher true discovery rates and lower FDR, with improvements amplified at larger sample sizes (Fig.2c). These results highlight the importance of modelling time-varying associations when performing feature selection in longitudinal settings.

### TPClust identifies cognitive subtypes with distinct trajectories and time-varying risk profiles

To evaluate whether TPClust can uncover clinically meaningful heterogeneity in cognitive aging, we applied the TPClust to 1,020 older adults from the ROSMAP (Bennett et al., 2018), two harmonized longitudinal cohorts of individuals without known dementia at baseline. Global cognition, derived from 19 neuropsychological tests, served as the longitudinal outcome. Covariates included sex, *APOEε*4 status, and longitudinal measurements of hypertension, diabetes, stroke, and frailty. Molecular predictors consisted of 2,015 differentially expressed genes (FDR < 0.05) from dorsolateral prefrontal cortex (DLPFC) tissue, annotated to 16 Gene Ontology biological pathways.

TPClust identified four subtypes with distinct age-related cognitive trajectories estimated from the model (Table 1; Fig. 3a; Fig. 3f). The largest group, *Resilient* (*n* = 642, 63%), exhibited stable cognitive function across the observed age range. *Late-Onset Decline* (*n* = 102, 10%) showed preserved cognition until age 85, followed by gradual decline. *Early Vulnerability* (*n* = 76, 7%) began at a lower cognitive baseline and declined from age 83.5. *Rapid Decline* (*n* = 200, 20%) exhibited accelerated decline starting before age 75. Estimated change points—defined as the ages at which the model detects a shift in cognitive trajectory—were 85.1, 88.1, 86.3, and 83.5 for the four subtypes, respectively (Fig. 3b). These values were broadly consistent with the 86.7-year change point reported by Yu et al. (2012) after which cognitive decline accelerated nearly fourfold in ROSMAP participants who were non-demented at baseline but later developed AD.

**Table 1:** Participant characteristics across TPClust-identified subtypes. P-values were calculated using ANOVA for continuous variables and the Chi-square test for categorical variables.

|  | *Resilient* | *Late-Onset Decline* | *Early Vulnerability* | *Rapid Decline* | P-value |
| --- | --- | --- | --- | --- | --- |
| Cohort size | 642 | 102 | 76 | 200 | - |
| Women *n*(%) | 423 (65%) | 67 (65%) | 48 (63%) | 153 (76%) | 0.03 |
| Age at visit (mean, SD) | 84.57 (7.25) | 84.30 (7.11) | 83.86 (7.06) | 84.94 (6.85) | 0.003 |
| Global cognitive function (mean, SD) | -0.04 (0.64) | -0.10 (0.72) | -0.48 (0.92) | -0.91 (1.16) | < 2E-16 |
| Pathological diagnosis of AD *n*(%) | 357 (56%) | 60 (59%) | 63 (83%) | 163 (81%) | <2E-12 |
| Diagnosis of AD dementia *n*(%) | 136 (27%) | 33 (45%) | 48 (86%) | 155 (96%) | < 2E-16 |
| *APOE*ε4 *n*(%) | 147 (23%) | 18 (18%) | 21 (28%) | 73 (36%) | 3.36E-04 |
| Diabetes *n*(%) | 125 (19%) | 12 (12%) | 18 (24%) | 29 (15%) | 0.17 |
| Hypertension *n*(%) | 617 (96%) | 98 (96%) | 74 (97%) | 192 (96%) | 0.96 |
| Stroke *n*(%) | 126 (20%) | 21 (21%) | 12 (16%) | 53 (26%) | 0.13 |
| Frailty *n*(%) | 273 (43%) | 36 (35%) | 40 (53%) | 133 (66%) | 3.60E-09 |

**Figure 3:**
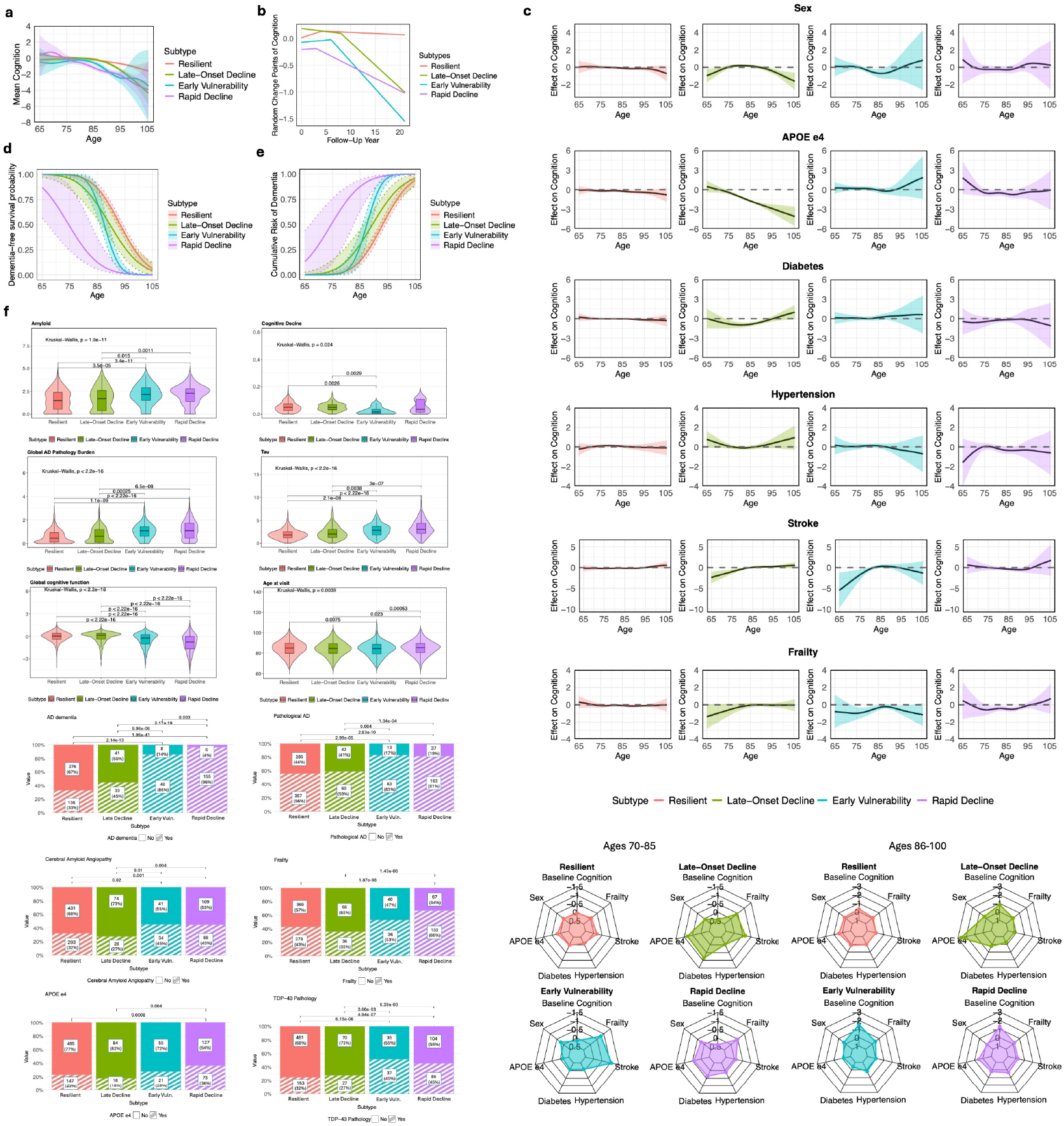
TPClust uncovers subtype-specific cognitive trajectories, time-varying covariate effects on cognition, and clinico-pathological distinctions. a, Estimated global cognitive trajectories with 95% confidence intervals across TPClust-identified subtypes. b, Distribution of individualized change points in global cognition across subtypes. c, Estimated time-varying effects of covariates (sex, *APOEε*4, diabetes, hypertension, stroke, frailty) on global cognition across age, shown by subtype. Radar plots summarize average effects of each covariate and baseline cognition, as estimated by the intercept function in TPClust, for two age groups (70–85 and 86–100) across subtypes. d, Dementia-free survival curves with 95% confidence intervals across subtypes. e, Cumulative incidence of Alzheimer’s disease with 95% confidence intervals by subtype. f, Distributions of clinico-pathological traits across subtypes.

To characterize subtype-specific dynamics of risk factors over age, we examined the time-varying covariate–cognition associations estimated by TPClust (Fig. 3c; Table S2). *Resilient* showed minimal associations, with only modest late-life effects of *APOEε*4 (ages 85–100), male (87–91), and stroke in narrow windows. *Late-Onset Decline* exhibited sustained vulnerability: *APOEε*4 (74–105), diabetes (74–92), and stroke (65–81) were associated with lower cognition, though stroke appeared protective at later ages (91–99). Male showed bidirectional effects—associated with lower cognition in early old age (65–70) and late old age (93–105), but higher cognition in mid old age (79–85). In *Early Vulnerability*, stroke (65–79) and male (79–89) were associated with lower cognition in early life, while frailty showed consistent effects across both early (72–86) and late old age (91–101) aging. *Rapid Decline* demonstrated broad susceptibility, with *APOEε*4 (76–90), frailty (76–93), and stroke (85–91) consistently associated with lower cognition.

To facilitate comparison, we averaged the estimated time-varying covariate effects across early (70–85) and late (86–100) age intervals (Fig. 3c). This revealed clear divergence across subtypes: early aging was largely unremarkable in *Resilient* but marked by emerging vulnerability in *Late-Onset Decline APOEε*4, diabetes, stroke, frailty), vascular and aging-related burden in *Early Vulnerability* (stroke, frailty), and multifactorial risk in *Rapid Decline* (male, *APOEε*4, diabetes, frailty). These patterns persisted in late life, with *Resilient* remaining largely unaffected, *Early Vulnerability* dominated by frailty, and *Rapid Decline* maintaining a multifactorial risk profile led by *APOEε*4 with modest contributions from diabetes, hypertension, stroke, and frailty.

To assess subtype-specific differences in progression to clinical dementia, we conducted dementia-free survival analyses across subtypes. Dementia-free survival differed significantly (log-rank *p <* 10^*−*3^), with median age at AD dementia onset estimated at 92.9, 89.5, 87.3, and 74.0 years for *Resilient, Late-Onset Decline, Early Vulnerability*, and *Rapid Decline*, respectively. At age 85, survival probabilities were highest in *Resilient* (84.6%) and lowest in *Rapid Decline* (11.9%), with intermediate rates in *Late-Onset Decline* (70.1%) and *Early Vulnerability* (70.5%) (Fig. 3d). Cumulative dementia incidence exceeded 87% in *Rapid Decline* by age 85, compared to ~29% in *Late-Onset Decline* and *Early Vulnerability*, and only 15% in *Resilient* (Fig. 3e; Tables S3–S4). After age 85, *Early Vulnerability* showed faster cognitive deterioration than *Late-Onset Decline*. After age 80, all non-*Resilient* subtypes showed steeper declines in dementia-free survival, highlighting distinct clinical progression trajectories.

### Multimodal profiling reveals clinico-pathological and molecular divergence across TPClust subtypes

To evaluate the biological and clinical relevance of TPClust-defined subtypes, we examined subtype-specific characteristics across clinical and neuropathological traits, molecular biomarkers, transcriptomics, proteomics, and neuroimaging.

We first examined clinical and neuropathological traits across subtypes (Table S5-S7; Fig. 4a–b). Compared to other subtypes, *Resilient* and *Late-Onset Decline* exhibited lower levels of *β*-amyloid deposition, tau tangle density, TDP-43, and cerebral amyloid angiopathy (CAA), alongside slower cognitive decline and higher residual cognition (De Jager et al., 2018; Yu et al., 2015) and Mini-Mental State Examination (MMSE) scores (Trzepacz et al., 2015). The *Early Vulnerability* showed intermediate levels of neuropathology and cognitive impairment. *Rapid Decline* showed the most severe clinico-pathological profile, with elevated *β*-amyloid, tau, TDP-43, and CAA, along with higher prevalence of AD dementia, pathological AD, *APOEε*4 carriers, atherosclerosis, hippocampal sclerosis, neocortical Lewy bodies, and dementia with Lewy bodies. This subtype also showed the fastest cognitive decline and the lowest residual cognition, MMSE scores, and total daily activity levels (James et al., 2012). Furthermore, the *Rapid Decline* subtype exhibited the highest prevalence of acute and subacute infarction (Arvanitakis et al., 2011) in both cortical and subcortical regions. To assess whether subtype differences in cognitive decline were explained by AD pathology, Braak stage, or frontal white matter integrity (R2), we repeated the analysis adjusting for these variables as covariates. Subtype effects remained significant (Table S8), and some contrasts—such as *Resilient* vs. *Late-Onset Decline*—became more pronounced after adjustment, suggesting that the observed differences in cognitive decline reflect biologically distinct and potentially uncharacterized mechanisms. Visualization using t-distributed stochastic neighbor embedding (t-SNE) of 618 individuals based on neuropathological traits revealed a smooth continuum from *Resilient* to *Rapid Decline* (Fig. 4f). Co-pathology burden also increased along this gradient: approximately 70% of *Early Vulnerability* and *Rapid Decline* had either TDP-43 or Lewy body pathology, compared to fewer than 40% in the *Resilient* and *Late-Onset Decline*.

**Figure 4:**
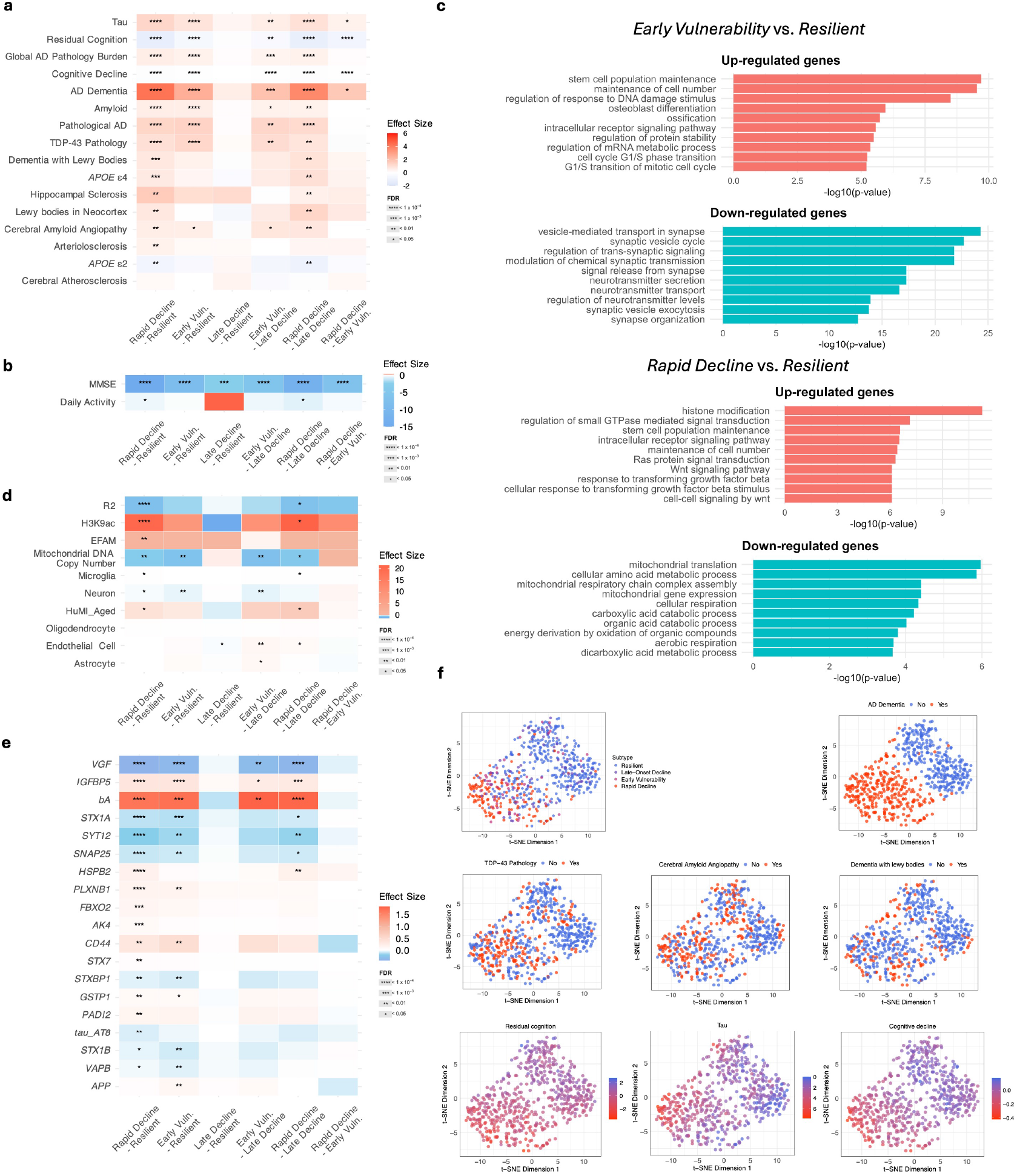
TPClust-derived subtypes exhibit distinct clinico-pathological, molecular, and transcriptomic signatures of Alzheimer’s disease. a, Associations between subtypes and clinico-pathological traits. b, Associations with MMSE scores and baseline daily activity. c, GO enrichment analysis for differentially expressed genes between *Early Vulnerability* vs *Resilient* and *Rapid Decline* vs *Resilient*, highlighting up- and down-regulated pathways. d, Associations with molecular and cellular biomarkers. e, Associations with targeted proteomics data via Selected Reaction Monitoring. f, t-SNE visualization of subtype membership and clinico-pathological traits, illustrating subtype separation and trait-specific distributions.

We next evaluated molecular and cellular biomarkers (Fig. 4d; Table S9). *Rapid Decline* showed elevated histone H3K9 acetylation (H3K9ac), the Epigenomic Factor of Activated Microglia (EFAM)—a CpG methylation-based index of microglial activation (Ma et al., 2022)—and increased expression of the HuMi-Aged gene set enriched in aged microglia (Olah et al., 2018), suggestive of age-related epigenetic changes and microglial activation. In contrast, *Resilient* exhibited preserved R2, higher neuronal proportions, and increased mitochondrial DNA copy number. *Early Vulnerability* showed reduced neuronal content and mitochondrial integrity compared to *Late-Onset Decline*, and increased endothelial cell proportions compared to *Resilient* and *Late-Onset Decline*.

Among 20 transcripts selected by TPClust (Table S10), the *Rapid Decline* was enriched for *PPM1G, PDZRN3, NPNT*, and *ADAMTS2*—genes implicated in tau signaling, blood–brain barrier integrity, and amyloid-related neurodegeneration (Gueniot et al., 2022; Felsky et al., 2023). In contrast, *RPH3A*, a synaptic resilience gene (Yu et al., 2023) was consistently downregulated across impaired subtypes. *MEIS3*, previously associated with lower Braak stage (Li and De Muynck, 2021), was most highly expressed in *Late-Onset Decline*, moderately elevated in *Early Vulnerability*, and suppressed in *Rapid Decline*. Transcriptome-wide analyses (Tables S11–S17; Fig. 4c) revealed distinct subtype-specific pathways. *Rapid Decline* showed upregulation of histone modification and Ras/Wnt signaling, and downregulation of mitochondrial respiration and amino acid metabolism. *Early Vulnerability* was enriched for DNA damage response and proteostasis pathways and showed reduced synaptic signaling.

Proteomics analysis (Fig. 4e; Table S18) showed *Resilient* had elevated synaptic proteins (*SNAP25, STX1A, STX1B, STXBP1, SYT12*, and *VGF*), previously implicated in synaptic maintenance in AD (Ramos-Miguel et al., 2021; Yu et al., 2023). In contrast, *Rapid Decline* showed increased *β*-amyloid (*bA*), glial and inflammatory markers (*CD44, PLXNB1*), and stress- and mitochondrial-related proteins (*GSTP1, AK4, HSPB2, FBXO2, IGFBP5*), which have been associated with neurodegeneration and cognitive decline in AD (Yu et al., 2018; Sullivan et al., 2019). *Early Vulnerability* displayed intermediate profiles, including elevated *bA* and *IGFBP5*, previously associated with synaptic loss and cellular stress (Miao et al., 2025).

Finally, neuroimaging data (Table S19) revealed significant cortical and subcortical atrophy in *Rapid Decline*, including the hippocampus and entorhinal cortex, while *Early Vulnerability* or *Late-Onset Decline* showed preserved brain structure.

### Integrative modeling in TPClust outperforms unimodal subtyping approaches

We benchmarked TPClust’s integrative model against unimodal subtyping methods based on either clinical or omics data alone to assess subtype resolution and their ability to delineate clinico-pathological heterogeneity.

For the clinical-only comparison, we applied TPClust to longitudinal cognitive trajectories in ROSMAP, excluding omics features but retaining all other model settings. This yielded two subtypes—*Resilient clinic* and *Rapid Decline clinic*—with *Resilient clinic* overlapping 80.0% with TPClust *Resilient*, and *Rapid Decline clinic* comprising individuals from *Rapid Decline* (76.8%) and *Early Vulnerability* (17.0%) (Fig. 5a). These subtypes recapitulated TPClust’s cognitive extremes but collapsed intermediate heterogeneity. Clinico-pathological contrasts mirrored those between TPClust *Resilient* and *Rapid Decline* but with reduced resolution (Fig. 5e; Table S20).

**Figure 5:**
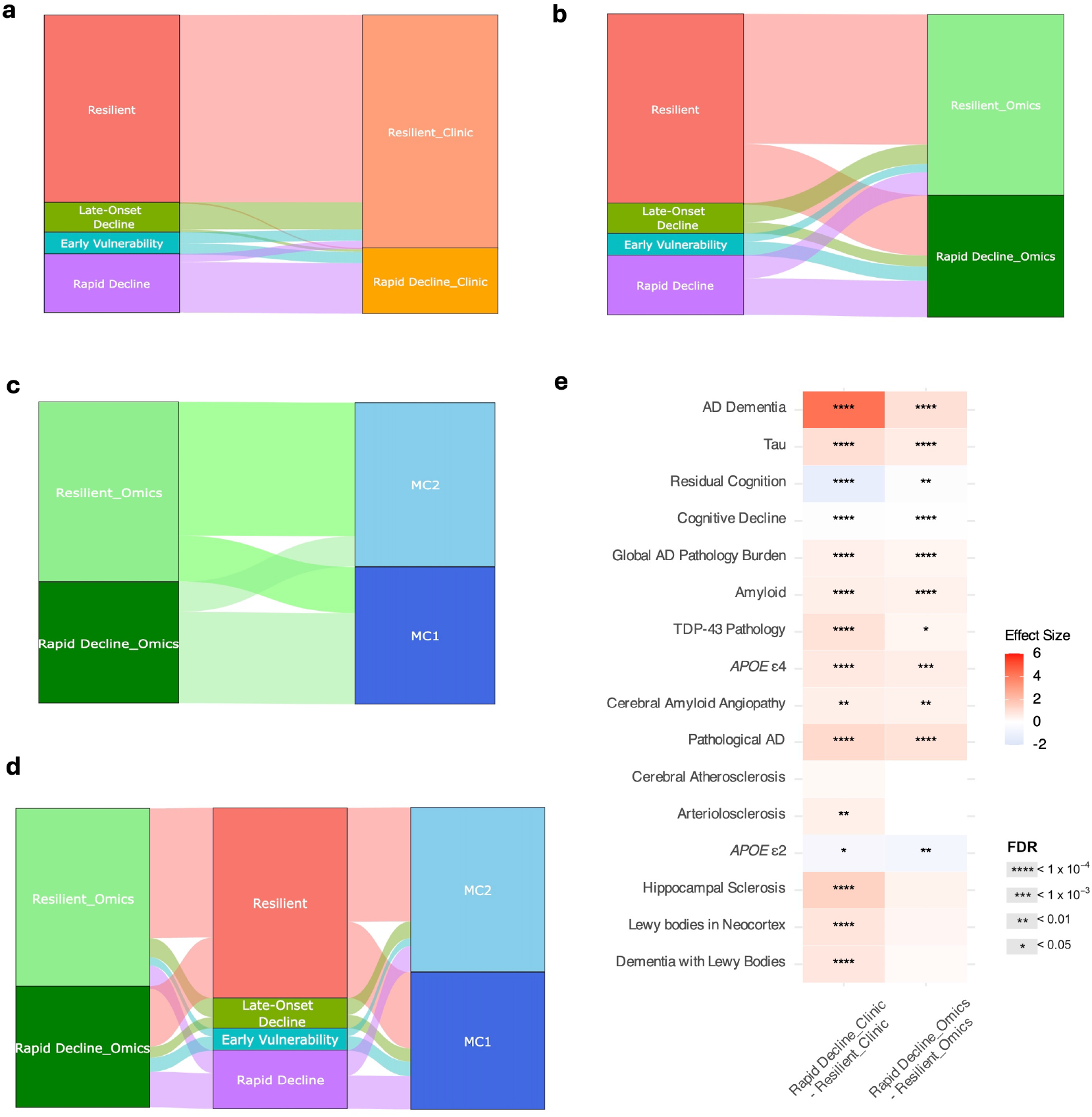
TPClust outperforms unimodal subtyping in capturing AD-related traits. a, Sankey diagram showing correspondence between TPClust-defined subtypes and subtypes derived from a clinical-only model. b, Correspondence between TPClust-defined subtypes and those from an omics-only model. c, Mapping between omics-only subtypes and transcriptome-based molecular clusters (MC1 and MC2). d, Integrated Sankey diagram linking omics-only, TPClust, and transcriptomic clusters, demonstrating improved subtype resolution and clinico-pathological specificity with TPClust. e, Associations of clinical-only and omics-only subtypes with clinico-pathological traits. Heatmap shows standardized effect sizes and FDR-adjusted significance levels.

For the transcriptomics-only comparison, principal component analysis (PCA) was performed on DLPFC gene expression data in ROSMAP (retaining 308 principal components explaining 90% of the variance), followed by K-means clustering with two clusters. The resulting *Resilient omics* subtype overlapped 71.9% with TPClust *Resilient*, while *Rapid Decline omics* was more heterogeneous, comprising individuals from *Rapid Decline* (29.7%), *Early Vulnerability* (11.7%), and *Resilient* (49.6%) (Fig. 5b). This broad distribution diminished clinico-pathological resolution and did not delineate subtypes with distinct pathological signatures, including arteriolosclerosis, hippocampal sclerosis, neocortical Lewy bodies, and dementia with Lewy bodies (Table S21). Additional transcriptomics-based comparisons are provided in the Supplementary Information (Fig. 5c-d).

## Discussion

Subtyping analysis is widely used to uncover population heterogeneity in complex diseases. Yet, existing approaches often rely on cross-sectional data, assume static covariate effects, and do not jointly model high-dimensional molecular features with longitudinal clinical outcomes. This can result in subtypes that are clinically and biologically ambiguous or lead to overconfident interpretations that obscure underlying disease mechanisms. To address this problem, we developed TP-Clust, a supervised, semi-parametric clustering method that integrates high-dimensional omics data with longitudinal phenotypes—including outcomes and covariates—for outcome-guided subtyping. TPClust jointly models subtype membership and outcome trajectories by combining a multinomial logistic model informed by omics features with spline-based regression for subtype-specific, time-varying covariate effects. Feature selection is performed using structured regularization that incorporates lasso, group lasso, and sparse group lasso penalties, allowing robust identification of biologically informative features while preserving model interpretability in high-dimensional settings.

In simulation studies, TPClust provided valid inference for time-varying covariate effects and robust feature selection in high-dimensional longitudinal data. Bootstrap-based confidence intervals achieved empirical coverage near the nominal 95% level, and trajectory estimation error decreased with increasing sample size, indicating improved estimation of subtype-specific time-varying covariate effects. In feature selection, TPClust consistently outperformed a high-dimensional mixture model with static covariate effects, achieving higher true discovery rates and lower false discovery rates.

Finally, we applied TPClust to transcriptomics and longitudinal cognitive data from the ROSMAP cohort, identifying four clinically and biologically distinct subtypes. *Resilient* individuals maintained stable cognition with preserved synaptic, neuronal, and mitochondrial integrity, minimal neuropathology, and weak risk factor associations, aside from modest late-life effects of *APOEε*4 and stroke. *Late-Onset Decline* showed delayed decline, moderate pathology, and sustained vulnerability to *APOEε*4, diabetes, and stroke. *Early Vulnerability* began with lower baseline cognition and early-life effects of stroke and frailty, with persistent frailty burden, reduced neuronal and mitochondrial content, synaptic loss, and elevated proteostasis stress. *Rapid Decline* displayed the most severe clinical and pathological profile, with early dementia onset and severe AD pathology—including *β*-amyloid, tau, TDP-43, CAA, atherosclerosis, hippocampal sclerosis, and Lewy bodies—alongside broad molecular disruption (e.g., epigenetic dysregulation, oxidative stress, glial activation, blood–brain barrier dysfunction, white matter degeneration), and multifactorial time-varying risk driven by *APOEε*4, frailty, and vascular factors. TPClust reveals clinically meaningful subtypes by modelling dynamic risk trajectories and integrating them with molecular profiles, offering a generalizable method for dissecting disease heterogeneity.

Compared to clinical- or transcriptomics-only subtyping approaches, TPClust identified more refined and biologically coherent subtypes, including intermediate profiles that were obscured by unimodal methods. Clinical-only modeling recovered cognitive extremes but collapsed intermediate heterogeneity, while omics-only clustering produced diffuse groupings with limited neuropathological resolution.

A limitation of TPClust is its current restriction to a single continuous outcome. Many clinical applications involve multiple longitudinal or categorical measures and extending the model to handle multivariate or mixed-type outcomes would improve its utility for multimodal phenotyping. Another limitation is the need to pre-specify the number of subtypes, typically chosen through cross-validation. While effective, this approach does not capture uncertainty in cluster number; model-based criteria or Bayesian nonparametric strategies could improve flexibility and robustness. Finally, TPClust is designed for retrospective analyses. Adapting the model for prospective prediction and early-stage stratification based on incomplete trajectories could enhance its clinical utility.

## Methods

### Model Specifications for TPClust

TPClust integrates high-dimensional omics data, longitudinal outcomes, and both longitudinal and cross-sectional covariates to infer biologically and clinically meaningful subtypes. The model is implemented in two steps. In the first step, TPClust estimates latent subtype memberships, selects informative omics features, and characterizes subtype-specific, time-varying covariate effects. In the second step, it constructs pointwise 1 *− α*-confidence intervals for these time-varying effects using a multiplier bootstrap procedure. Unlike conventional bootstrap methods that rely on resampling observations, the multiplier bootstrap generates independent random weights to individual likelihood contributions, resulting in a weighted objective function for parameter estimation. By repeating this procedure multiple times, TPClust approximates the distribution of time-varying effects and construct confidence intervals based on a normal approximation to the bootstrap distribution.

Let ***g***_*i*_ = (1, *g*_*i*1_, …, *g*_*iQ*_)^*τ*^ denote the high-dimensional omics features for subject *i*, including an intercept term, where *Q* is the number of omics features. Each subject may contribute multiple longitudinal observations indexed by visit time *t*_*ij*_, representing the time of individual *i* at their *j*-th visit —for example, age or time since baseline. Let *y*_*i*_(*t*) denote the smooth outcome trajectory for subject *i* with *y*_*i*_(*t*_*ij*_) representing the observed outcome at each visit, and let ***y***_*i*_ denote the corresponding vector of outcome values across visits.

#### Supervised, semi-parametric clustering for outcome-guided subtyping using longtudinal clinical data and high-dimesional omics data

To identify clinically meaningful subtypes, we adopt a supervised mixture regression framework that models subtype-specific associations between longitudinal outcomes and covariates. Let *f*_*k*_(***y***_*i*_; ***x***_*i*_, ***v***_*i*_) denote the outcome model for subtype *k, k* = 1, …, *K*, and *K* is the total number of subtypes. Since the subtype assignment is unobserved, we define the overall likelihood for individual *i* by marginalizing over subtype membership using a mixture of subtype-specific models:

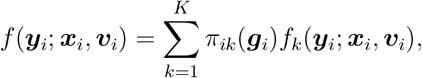

where *π*_*ik*_(***g***_*i*_) denotes the probability that individual *i* belongs to subtype k, modeled as a function of the high-dimensional omics features ***g***_*i*_, and constrained such that 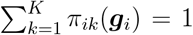, for all *i* = 1, …, *n*. Further details on the specification of *f*_*k*_(***y***_*i*_; ***x***_*i*_, ***v***_*i*_) and *π*_*ik*_(***g***_*i*_) are provided below.

#### Omics-based sub-model for outcome-guided subtyping

High-dimensional Omics profiles are often informative for identifying subtypes that reflect underlying biological heterogeneity, particularly in complex disorders such as neurodegenerative diseases. Motivated by prior work using unsupervised clustering on omics data (Milind et al., 2020; Neff et al., 2021; Lee et al., 2023; Sahelijo et al., 2024), our framework incorporates molecular information into subtype assignment to capture clinically relevant heterogeneity. Let ***z***_*i*_ = (*z*_*i*1_, …, *z*_*iK*_) denote the latent subtype labels for subject *i*, where *z*_*ik*_ = 1 if individual *i* belongs to subtype *k* and 0 otherwise. We model the probability of subtype membership as a function of the omics profile ***g***_*i*_ via a multinomial logistic model:

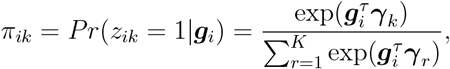

where ***γ***_*k*_ = (*γ*_*k*0_, *γ*_*k*1_, …, *γ*_*kQ*_) are the subtype-specific regression parameters. To ensure model identifiability, we fix ***γ***_1_ = **0**.

#### Longitudinal sub-model for time-varying clinical associations

In many progressive diseases, clinical manifestations and risk factors effects evolve with time. For example, the effects of *APOE ε*4 and early-onset diabetes on AD risk vary with age (Tang et al., 1998; Bellou et al., 2020; Amidei et al., 2021). Similarly, in Parkinson’s disease (PD), diabetes severity was shown to interact with age, with greater diabetes severity associated with increased hazard of PD onset among individuals aged 40–65 (Han et al., 2023). To flexibly capture subtype-specific, time-varying effects of both static and dynamic covariates on longitudinal outcomes, we model the outcome *y*_*i*_(*t*_*ij*_) for subject *i* at time *t*_*ij*_, conditional on subtype *k*, as:

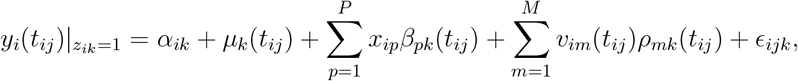

where *α*_*ik*_ is a random effect capturing the covariance of longitudinal outcomes associated with the same individual with 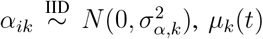is the mean outcome trajectory for subtype *k, β*_*pk*_(*t*)’s and *ρ*_*mk*_(*t*)’s are smooth time-varying effects for cross-sectional covariate *x*_*ip*_ and longitudinal covariate *v*_*im*_ (*t*), respectively, and *ϵ* _*ijk*_ represents the random noise term with 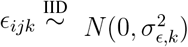. To notational simplicity, we set *x*_*i*0_ = 1 for all *i* = 1, …, *n* and use *β*_0*k*_(*t*) to denote *µ*_*k*_(*t*), allowing the model to be compactly rewritten as:

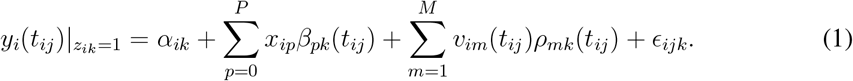

### Model Implementation

To estimate the subtype-specific time-varying covariate effects, *β*_*pk*_(*t*) and *ρ*_*mk*_(*t*), we approximate each unknown functions using finite-dimensional non-parametric B-spline basis expansions. Based on these approximations, we formulate a likelihood-based optimization problem, which we solve using a modified iterative expectation-maximization (EM) algorithm. This procedure enables simultaneous estimation of time-varying covariate effects and selection of informative features in high-dimensional omics data.

#### Nonparametric spline approximation for time-varying associations

To flexibly model time-varying covariate effects, we approximate each smooth function using a finite-dimensional B-spline basis expansion (De Boor, 1986). Specifically, we represent the subtype-specific time-varying effects as:

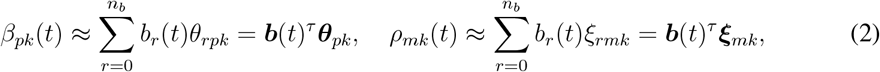

where 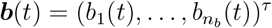 denotes the vector of B-spline basis functions, and ***θ***_*pk*_ and ***ξ***_*mk*_ are the basis coefficients for covariates *x*_*ip*_ and *v*_*im*_, respectively, in subtype *k*. With (2), we can approximate (1) by

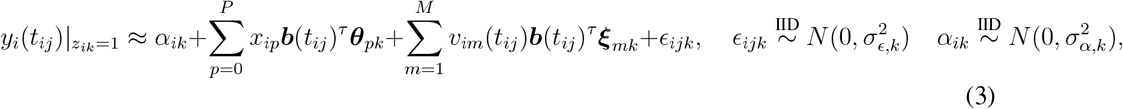

where 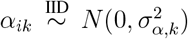 is the subject-specific random intercept and 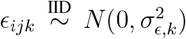 measurement error.

#### Likelihoods

Let 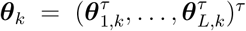 and 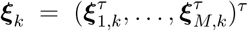. Define the complete parameter vector as ***η*** = *{****θ, ξ, γ, σ****}*, where ***θ*** = (***θ***_1_, …, ***θ***_*K*_)^*τ*^, ***ξ*** = (***ξ***_1_, …, ***ξ***_*K*_)^*τ*^, ***γ*** = (***γ***_1_, …, ***γ***_*K*_)^*τ*^ and ***σ*** = (*σ*_*ϵ*,1_, …, *σ*_*ϵ,K*_, …, *σ*_*α,K*_)^*τ*^. The observed data likelihood is given by

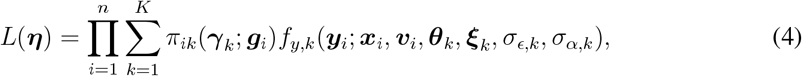

where *π*_*ik*_(***γ***_*k*_; ***g***_*i*_) is the subtype membership probability and *f*_*y,k*_(***y***_*i*_; ***x***_*i*_, ***v***_*i*_, ***θ***_*k*_, ***ξ***_*k*_, *σ*_*ϵ,k*_, *σ*_*α,k*_) is the likelihood under the model (3).

#### Expectation-maximization algorithm and likelihood-based estimation

Direct maximization of the observed-data log-likelihood (4), is computationally challenging. To address this, we adopt a modified EM algorithm by treating the latent subtype labels ***z***_*i*_ and random effects ***α***_*i*_ = (*α*_*i*1_, …, *α*_*iK*_) as unobserved variables and derive the likelihood conditional on ***z***_*i*_ and ***α***_*i*_. This leads to an optimization problem that is more tractable to solve. Specifically, we define the complete data as 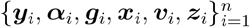, and treat ***α***_*i*_ and ***z***_*i*_ as missing since they are not directly observed. We use *T*_*i*_ to denote the total number of visits for individual *i*. Let *f*_*y,k*|***α***_(*y*_*i*_(*t*_*ij*_); *α*_*ik*_, ***x***_*i*_, ***v***_*i*_, ***θ***_*k*_, ***ξ***_*k*_, *σ*_*ϵ,k*_) denote the likelihood of the clinical outcome *y*_*i*_(*t*_*ij*_) conditional on ***z***_*i*_ and ***α***_*i*_, and let *f*_*α,k*_(*α*_*ik*_; *σ*_*α,k*_) denote the density of the random effect *α*_*ik*_. Then, the complete-data log-likelihood for ***η*** can be written as:

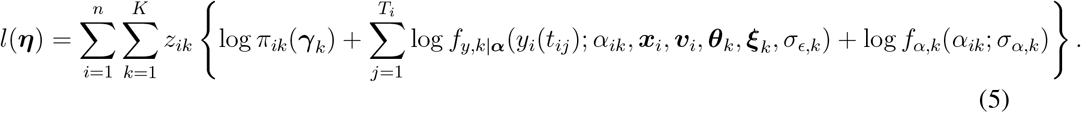

The log-likelihood (5) can be decomposed into two components:

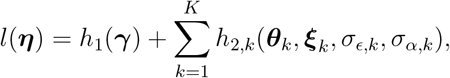

where

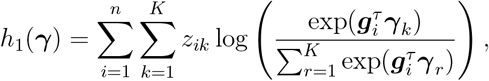

represents the contribution of the multinomial logistic submodel for omics data, and

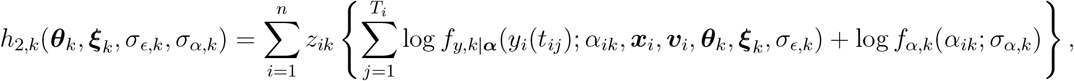

reflects the contribution from the longitudinal outcome model for subtype *k*. According to the above decomposition, the maximization of *l*(***η***) is equivalent to the maximization of *h*_1_(***γ***) and *h*_2,*k*_(***θ***_*k*_, ***ξ***_*k*_, *σ*_*ϵ,k*_, *σ*_*α,k*_) for each *k* = 1, …, *K* independently. Using the method of profiling, the maximization of *h*_2,*k*_(***θ***_*k*_, ***ξ***_*k*_, *σ*_*ϵ,k*_, *σ*_*α,k*_) can be divided into two steps. In the first step, we estimate ***θ***_*k*_ and ***ξ***_*k*_ by maximizing the following objective function:

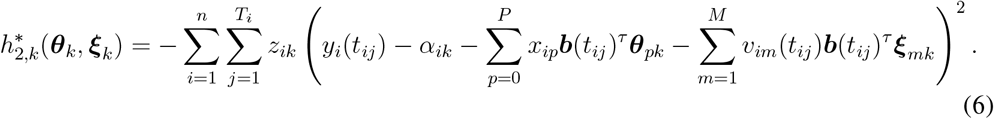

Let 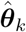 and 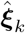 denote the maximizer of 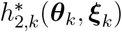. Given 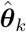 and 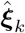, the residual variances *σ*_*ϵ,k*_ and *σ*_*α,k*_ are then updated via closed-form estimators:

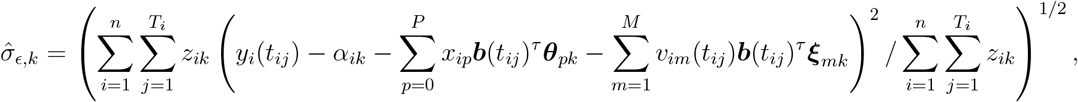

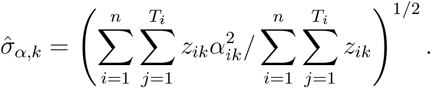

Define the profiled log-likelihood as:

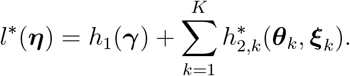

Then resulting profiled log-likelihood *l*^*\**^(***η***) shares the same maximizer as the full log-likelihood *l*(***η***), ensuring valid and consistent parameter estimation within the EM framework.

#### Feature selection for high-dimensional omics

High-throughput omics data are typically high-dimensional, and incorporating irrelevant features can introduce bias and reduce interpretability. To address this, our framework integrates a biologically informed feature selection mechanism during parameter estimation. Several regularization techniques have been proposed for high-dimensional variable selection, including LASSO (Tibshirani, 1996), elastic net (Zou and Hastie, 2005), and sparse group LASSO(Simon et al., 2013).

We adopt a sparse group LASSO regularization framework that combines two complementary penalties: a group LASSO penalty to encourage sparsity at the biological pathway level, and a standard LASSO penalty to induce sparsity among individual omics features. Omics features are grouped based on prior biological annotations, allowing the model to eliminate entire non-informative groups while retaining informative features within selected pathways. In parallel, the standard LASSO component enables discovery of relevant features not assigned to any pre-defined group. This hierarchical penalty structure improves estimation efficiency and biological interpretability by leveraging pathway-level prior knowledge, enabling the model to first eliminate entire irrelevant pathways and then select informative omics features within the retained pathways.

We denote *L* by the total number of known pathways, and by *A*_*l*_ the index set of omics associated with the pathway *l, l* = 1, …, *L*. Let 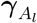 represent the subset of ***γ*** corresponding to the pathway *l*,

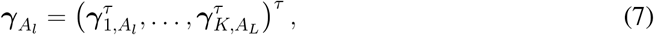

where 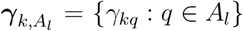, *k* = 1, …, *K*. To enable both feature- and pathway-level selection, we impose a sparse group LASSO penalty on gamma of the form:

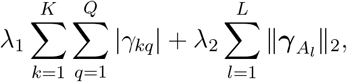

where *λ*_1_ and *λ*_2_ are regularization parameters controlling sparsity at the individual and group (pathway) levels, respectively. The first term promotes sparsity among all individual omics features, while the second term encourages group-wise shrinkage, effectively removing entire pathways when they are not informative.

#### Smooth estimation of time-varying covariate effects

To estimate the unknown time-varying covariate effects in our model, we approximate each function using B-spline basis expansions. The number of basis functions, denoted by *n*_*b*_, is typically set large to reduce approximation bias. However, a large *n*_*b*_ can lead to an ill-conditioned design matrix, resulting in numerical instability during optimization. Furthermore, even when a numerical algorithm successfully identifies an optimizer, the resulting estimates of the time-varying coefficients can be overly variable and lack smoothness. To address these challenges, we impose a roughness penalty on the spline coefficients to enforce smoothness of each estimated time-varying effect. Specifically, for the spline-based approximations of *β*_*pk*_(*t*) and *ρ*_*jk*_(*t*) in (2), we apply the following penalty to ***θ***_*pk*_ (with a similar form for ***ξ***_*jk*_):

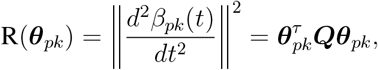

where ***Q*** is a symmetric and positive semi-definite matrix. This form of penalty, commonly known as a roughness penalty, is widely used in the functional data analysis literature to encourage smoothness in estimated curves (Ramsay and Silverman, 2005).

#### Penalized Likelihood for Joint Estimation and Selection

Finally, after incorporating omics feature selection and controlling the smoothness of the estimated time-varying covariate effects, we define the following penalized objective function:

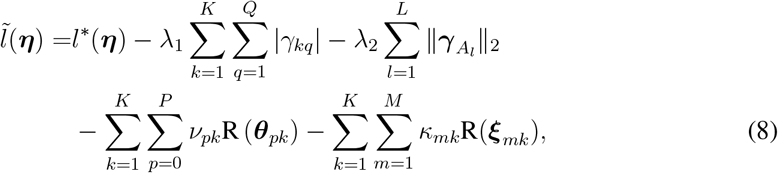

Here, *λ*_1_, *λ*_2_, *ν*_*pk*_, and *κ*_*mk*_ are tuning parameters that control the strength of the sparsity and smoothness penalties. In (8), the roughness penalties involve *K ×* (*P* + *M* + 1) tuning parameters, which can impose a substantial computational burden. To mitigate this issue, we simplify the structure by using a common tuning parameter *ν*_*p*_ for the roughness penalties applied to all *β*_*pk*_(*t*) across subtypes. Similarly, a common parameter *κ*_*m*_ is used for all *ρ*_*mk*_(*t*) across subtypes.This reduction in the number of tuning parameters improves computational efficiency while maintaining flexibility in the model. This leads to the following simplified penalized objective function:

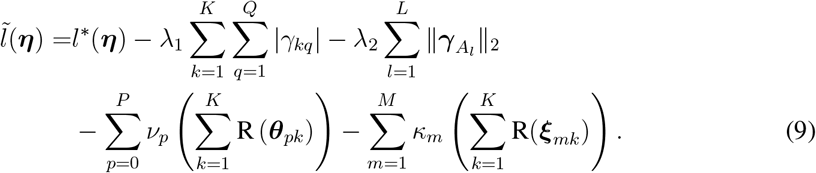

### Algorithm for Model Fitting

We estimate model parameters by maximizing the penalized objective function in Equation (9). To efficiently solve this optimization problem, we implement a modified Expectation-Maximization (EM) algorithm that iteratively updates parameter estimates until convergence.

#### The modified EM algorithm

Let *s* denote the iteration index. At each iteration, the algorithm alternates between the following steps:

The E-step at the *s*-th iteration computes the conditional expectation of the penalized objective function 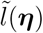 with respect to the latent variables ***z***_*i*_ and ***α***_*i*_, given the observed data ***y***_*i*_, ***x***_*i*_, and ***v***_*i*_, and the current parameter estimates ***η***^(*s*)^:

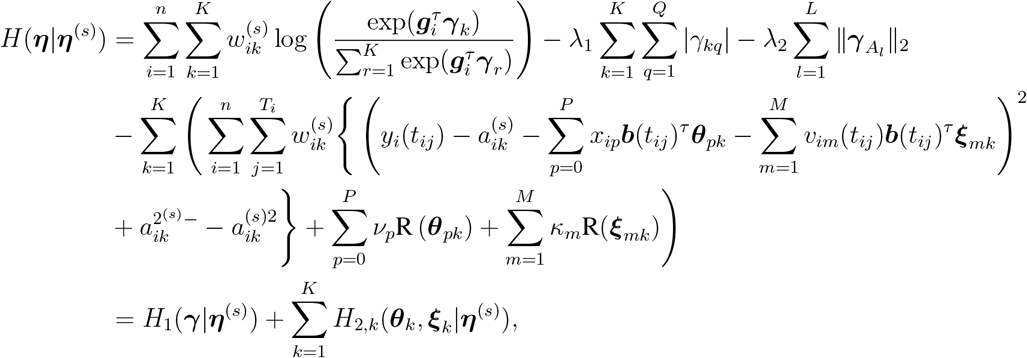

where

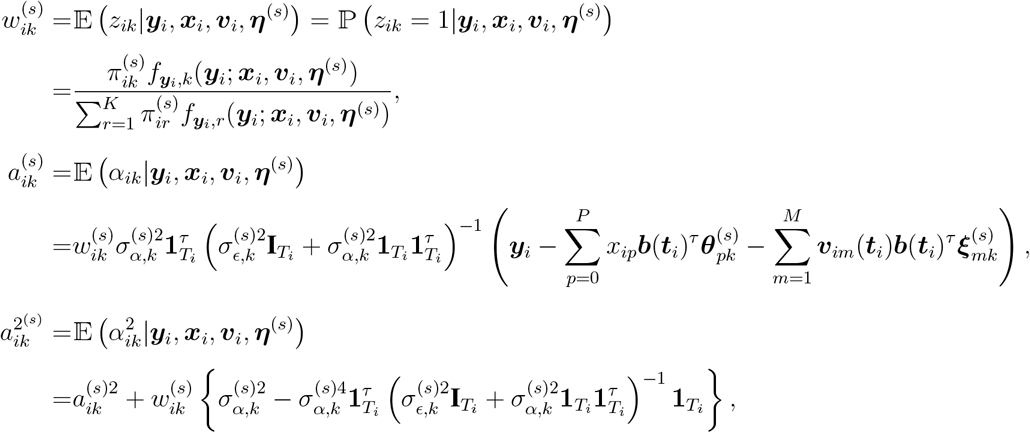

where *f*_***y***_*i*_,*k*_ denotes the joint density function of ***y***_*i*_.

The M-step at the (*s* + 1)-th iteration, maximizes *H*(***η*** | ***η***^(*s*)^) with respect to ***η***. Since *H*_1_(***γ***|***η***^(*s*)^) and *H*_2,*k*_(***θ***_*k*_, ***ξ***_*k*_|***η***^(*s*)^), for *k* = 1, …, *K* involve disjoint sets of parameters, the maximization of *H*(***η***|***η***^(*s*)^) can be decomposed into independent subproblems: maximizing *H*_1_(***γ***|***η***^(*s*)^) with respect to ***γ***, and maximizing each *H*_2,*k*_(***θ***_*k*_, ***ξ***_*k*_|***η***^(*s*)^) with respect to (***θ***_*k*_, ***ξ***_*k*_), for *k* = 1, …, *K*.

Maximization of *H*_1_:

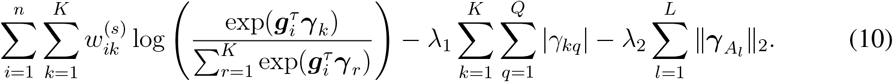

We propose to use disciplined convex programming (DCP) tools (Grant et al., 2009) to solve the maximization of (10). The direct maximization of (10) does not follow DCP rule sets. To overcome this, we convert the maximization of (10) into the maximization of the following objective function,

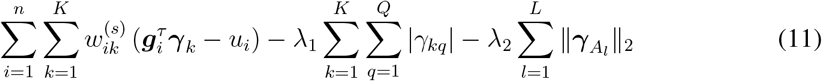

such that

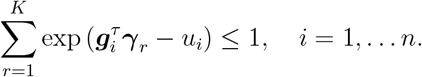

(10) and (11) share the same maximizer. The maximization problem in (11) adheres to the DCP rule set and can be efficiently solved using the R package CVXR with solver MOSEK. Additionally, the sparse group LASSO regularization in (11) can be replaced with alternative penalties, such as LASSO, group LASSO, or elastic net. These formulations can also be converted into a form compliant with the DCP rule set.

Maximization of *H*_2,*k*_:

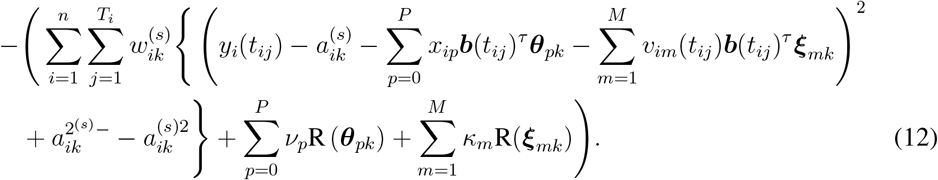

For a fixed *k*, we first vectorize the longitudinal outcomes *{y*_*i*_(*t*_*ij*_): *i* = 1, …, *n, j* = 1, …, *T*_*i*_*}* into a single column vector denoted by ***Y***. Next, we construct the design matrix ***X*** corresponding to the covariates *x*_*ip*_ and *v*_*im*_(*t*) after applying B-spline basis expansions—specifically, the terms *x*_*ip*_***b***(*t*_*ij*_)^*τ*^ and *v*_*im*_(*t*_*ij*_)***b***(*t*_*ij*_)^*τ*^ in (12). Let 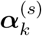 denote the vector of random effects 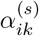 for *i* = 1, …, *n* at the *s*-th iteration, and let ***W***_*k*_ denote the diagonal matrix whose entries are the weights 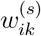 (expanded appropriately to match the dimensions of ***Y***). We also define ***U*** as a block-diagonal penalty matrix constructed from the tuning parameters *ν*_1_, …, *ν*_*P*_ and *κ*_1_, …, *κ*_*M*_. The maximizer of *H*_2,*k*_ at the (*s* + 1)-th iteration is then given by:

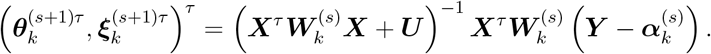

The detailed definition of ***Y***, ***X***, 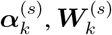 and ***U*** can be found in the supplementary document. Given 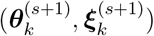`, we further update *σ*_*α,k*_ and *σ*_*ϵ,k*_ by

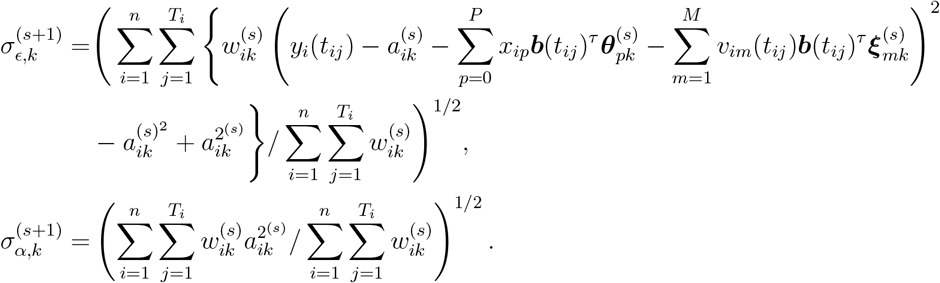

#### Parameter tuning strategy

The modified EM algorithm estimates model parameters for any given set of tuning parameters *λ*_1_, *λ*_2_, *ν*_1_, …, *ν*_*P*_, and *κ*_1_, …, *κ*_*M*_. To select these parameters, we adopt a strategy that combines the idea of relaxed LASSO (Meinshausen, 2007) with cross-validation. For any given set of tuning parameter values, we first fit the model using the entire dataset by maximizing the penalized objective function (9), which includes sparsity penalties. This yields an estimated set of informative omics features. Based on this, we partition ***γ***_*k*_ for each subtype *k* into two components: ***γ***_*k,S*_ and ***γ***_*k,N*_, where *S* denotes the index set of informative omics and *N* denotes the index set of non-informative omics. Next, we evaluate the quality of the selected omics using *K*-fold cross-validation (typically 5- or 10-fold). In each round of cross-validation, we follow the relaxed LASSO strategy and refit the model on the training data using only the informative features indexed by *S*. This refitting step excludes all sparsity penalties and focuses solely on the selected omics to estimate the model parameters—specifically, 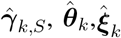, and 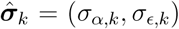 for *k* = 1, …, *K*. The entries of 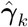 corresponding to the informative features are set to 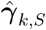, while those corresponding to non-informative features 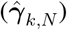 are fixed at zero. By eliminating the sparsity penalties in this step, we reduce the shrinkage bias introduced by LASSO-type regularization in the estimation of ***γ***_*k,S*_, *k* = 1, …, *K*. Using the refitted model, we compute the negative log-likelihood on the held-out test set as the validation loss. The optimal set of tuning parameters is then selected by minimizing this cross-validated loss.

### Multiplier Bootstrap-Based Inference

To conduct inference on the unknown time-varying effects *β*_*pk*_(*t*) and *ρ*_*mk*_(*t*), we adopt a multiplier bootstrap procedure to construct pointwise (1 *− α*) confidence intervals, where *α* denotes the significance level. Let *S* denote the index set of signal genes identified in the feature selection step, and let ***g***_*i,S*_ represent the signal gene measurements for individual *i* based on the index set *S*.

Let *b* denote the index of the bootstrap samples. For each bootstrap sample, we first draw *n* independent random weights 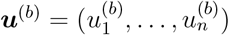 from the Rademacher distribution (i.e., each 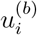 is independently sampled from {−1, 1} with equal probability). Using these weights, we define the following bootstrap analog of the log-likelihood components:

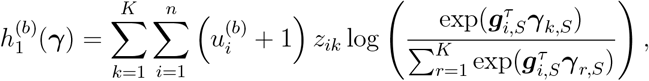

and

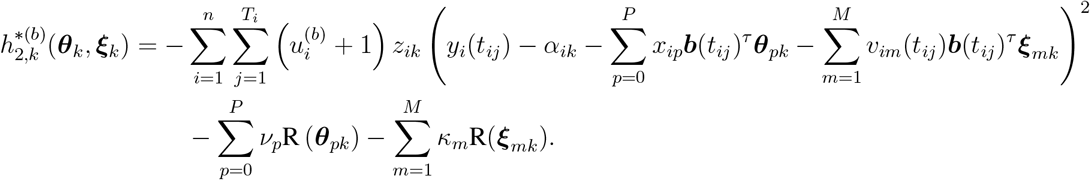

For each bootstrap sample, we estimate *β*_*pk*_(*t*) and *ρ*_*mk*_(*t*) by maximizing the following penalized objective function:

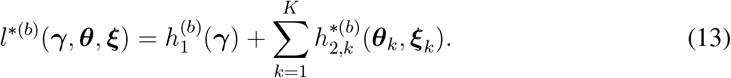

Let 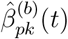 and 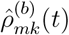 denote the bootstrap estimates obtained from the *b*-th bootstrap sample. Repeating this procedure *B* times yields the collection 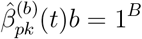, from which we compute the pointwise standard deviation, denoted by 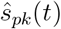. Under a normal approximation, the (1 *− α*) two-sided confidence interval for *βpk*(*t*) is given by:

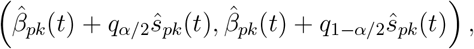

where *q*_*α/*2_ is the *α/*2 quantile of a standard normal. The confidence interval for *ρ*_*mk*_(*t*) can be constructed similarly.

### Simulated Study

#### Data generating models

We conduct two simulations to evaluate the performance of the TP-Clust method in terms of omics feature selection and estimation of time-varying associations among the longitudinal and cross-sectional variables. In both settings, the data are generated from a model with *K* = 3 subgroups, *P* = 2 cross-sectional covariates, and *M* = 1 longitudinal covariates.

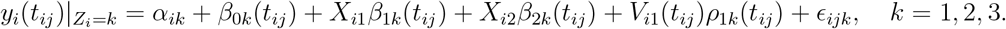

The mean curves *β*_0*k*_(*t*), and time-varying coefficients *β*_1*k*_(*t*) and *β*_2*k*_(*t*), are defined as linear combinations of B-splines. The time-varying coefficients *ρ*_*k*_(*t*) are either linear or quadratic functions. More details on the definition of these time-varying coefficients and data-generating procedure can be found in the supplementary document. In the first scenario, data are simulated based on *Q* = 110 omics features, among which 10 are informative. These features are grouped into *L* = 6 distinct pathways: one pathway contains all 10 informative features, while each of the remaining five pathways consists exclusively of 20 non-informative features. In the second scenario, the data are generated based on *Q* = 520 omics features organized into *L* = 7 pathways, with a total of 20 informative features. Two of the pathways each contain 10 informative features, and the remaining five pathways consist of 100 non-informative features each.

#### Metrics for evaluating TPClust in simulation studies

To evaluate the performance of the proposed estimators for the time-varying covariate effects *β*_*pk*_(*t*) and *ρ*_*mk*_(*t*), we use the integrated mean squared error (IMSE). Let *N*_rep_ denote the number of simulation repetitions. The IMSE of 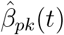 (or 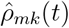 across *N*_rep_ repetitions is defined as:

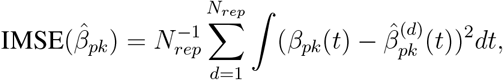

where 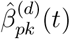 denotes the estimate obtained in the *d*-th simulation replication. A similar expression is used for computing IMSE 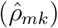. To evaluate the validity of the proposed multiplier bootstrap-based inference procedure, we examined the average coverage probabilities of each time-varying coefficient across 100 equally spaced time points in the interval [0, 1]. The nominal coverage level was set at 95%. Regarding the accuracy of omics feature selection, we assessed model performance using the false discovery rate (FDR) and the true discovery rate (TDR). The FDR is defined as the proportion of false positives among all selected omics features. For example, if 20 features are selected by the method as informative and 4 of them are actually non-informative, then the FDR is 4*/*20 = 0.2. The TDR is defined as the proportion of true positives among all true informative features. For instance, if there are 20 informative features and 18 of them are correctly selected, then the TDR is 18*/*20 = 0.9. We implemented the ogClust method using R function ogClust_GM from the package available at https://github.com/wenjiaking/ogClust.

### Study Design

#### Participants

The Religious Orders Study (ROS) and the Rush Memory and Aging Project (MAP) are two harmonized, prospective, community-based longitudinal cohort studies of aging (Bennett et al., 2018). Both cohorts enroll non-demented older adults at baseline, conduct annual clinical and neuropsychological assessments, and require participants to consent to brain donation at death. This design enables the integration of longitudinal cognitive trajectories with postmortem molecular and neuropathological data. At the time of death, participants span the full cognitive spectrum—from normal cognition to mild impairment to dementia—allowing disease progression to be modeled as a continuous process. For this study, molecular data consisted of bulk RNA-sequencing from dorsolateral prefrontal cortex (DLPFC) tissue obtained at autopsy. Following rigorous quality control and preprocessing, transcriptomic profiles were quantified for 16,674 unique genes (18,629 transcripts) across 1,092 participants.

#### Cognitive function

Global cognitive function was assessed at each visit using a composite z-score derived from a battery of 19 tests spanning five cognitive domain: episodic memory, semantic memory, working memory, perceptual orientation, and perceptual speed (Wilson et al., 2015). Raw scores from each domain were standardized to z-scores using the baseline mean and standard deviation of all participants, then averaged to create the composite global cognitive function score (Wilson et al., 2015).

#### Cardio-cerebrovascular risk factors

Cardio-cerebrovascular risk factors (vascular risk factors) were assessed at each visit based on self-reported medical history, self-reported use of disease-specific medications, and physical examinations (Lee et al., 2023). Hypertension was defined based on self-reported information on hypertension, systolic blood pressure *≥* 140 mmHg, diastolic blood pressure *≥* 90 mmHg, or use of anti-hypertension medications. Diabetes was defined using self-reported information on diabetes, hemoglobin A1C levels *≥* 7%, or use of diabetes medications. Stroke was ascertained through self-report and clinical diagnoses made by clinicians. Frailty was defined as the presence of three or more of the following five criteria: body mass index, fatigue, gait, grip strength, and physical activity (Buchman et al., 2007).

#### RNA sequencing data and processing

Gene-level transcription profiles were derived from the dorsolateral prefrontal cortex (DLPFC) tissue collected postmortem in ROSMAP. RNA-seq data were generated across multiple batches from various sequencing centers and experimental protocols, as previously described in Lee et al. (2023). Raw count data were normalized using the Trimmed Means of M-values method to create a frozen dataset available on Synapse (Synapse: syn25741873). Outlier samples were excluded based on expression profiles, and lowly expressed genes with a median raw count below 10 were filtered out to reduce technical noise. Covariate selection for expression modeling followed a forward selection approach (Lee et al., 2023). For DLPFC, selected covariates included age at death, sex, and technical confounding factors such as batch, library size, percentages of coding bases, aligned reads, ribosomal bases, UTR bases, and intergenic bases, as well as percentage duplication, median 5 prime to 3 prime bias, median 3 prime bias, median CV coverage, postmortem interval, and study index (ROS or MAP). To adjust for biological and technical confounding, linear regression was applied to model log 2-transformed normalized expression values as a function of these covariates. The resulting residuals —adjusted for demographic, study-specific, and sequencing-related variation— were used as the final transcription dataset. To account for transcript-level variability independent of demographic and technical factors, we used residualized gene expression values. Among the 18,629 transcripts profiled in the DLPFC, we identified 2,015 transcripts that were significantly differentially expressed between individuals with pathological AD and controls (FDR < 0.05; Table S22), providing a high-dimensional feature set enriched for AD-relevant biology. These residualized expression values of these 2,015 transcripts were used as input features for TPClust, with standard LASSO applied to all transcripts. In parallel, we performed Gene Ontology Biological Process (GO BP) enrichment analysis on the 2,015 differentially expressed transcripts, identifying 16 significantly enriched pathways (FDR < 0.05; Table S23). Transcripts annotated to these enriched pathways were then grouped to define input sets for the group LASSO penalty.

#### Proteomics data

Targeted proteomics data were obtained from the Selected Reaction Monitoring (SRM) assay in ROSMAP (Yu et al., 2018). The abundance of endogenous peptides was quantified as the log2 ratio relative to spiked-in heavy-isotope-labeled synthetic peptides. For normalization, these light/heavy log 2 ratios were median-centered within each sample to ensure a zero median. As a quality control measure, small aliquots from each homogenized sample were pooled and distributed throughout the study (8 per 96-well plate) to serve as QC samples. A signal-to-noise ratio was calculated as the variance across human subject samples divided by the variance across controls; peptides were excluded if their variance in controls was equal to or greater than that in the subject samples. Proteins measured by multiple peptides were summarized into a single protein-level value.

#### Neuroimaging data

We used structural MRI data from ROSMAP (Fleischman et al., 2025; Heywood et al., 2022). Participants underwent biennial imaging using multiple pulse sequences to capture structural, functional, and chemical brain characteristics. A standardized three-stage quality control pipeline was applied to ensure accuracy and reliability. First, phantom scans were used to assess scanner performance. Next, the quality of raw images was evaluated using both automated metrics and visual inspection. Finally, the derived imaging outputs were assessed for artifacts and overall accuracy. A total of 166 imaging-derived phenotypes were analyzed, including intracranial volume, regional gray matter volume, cerebrospinal fluid volume, white matter volume, white matter hyperintensities, regional cortical thickness measures, brain age, ARTS score, regional transverse relaxation rate (R2) in frontal white matter and other quantitative biomarkers relevant to neurodegeneration.

### Statistical Analysis

#### Using TPClust to identify clinically meaningful cognitive subtypes

To evaluate whether TP-Clust can identify clinically meaningful heterogeneity in cognitive aging, we applied the TPClust to 1,020 participants from the ROSMAP after excluding individuals with missing data. Global cognition served as the longitudinal outcome. Covariates included sex, *APOE ε*4 carrier status, and longitudinal measurements of hypertension, diabetes, stroke, and frailty. As the omics input for TPClust, we used the 2,015 differentially expressed genes (FDR < 0.05) derived from DLPFC tissue. TPClust incorporated this pathway structure to guide biologically informed gene selection during subtype discovery.

Once the TPClust model parameters were estimated, posterior probabilities of subtype membership were computed for each individual using the formula:

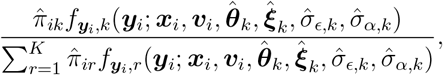

where 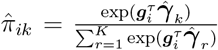 represents the estimated prior probability of individual *i* belonging to subtype *k* based on their omics features ***g***_*i*_. Each individual was assigned to the subtype corresponding to the highest posterior probability.

To examine the age-dependent associations between global cognitive function and clinical risk factors or confounders, we summarized the estimated time-varying covariate effects over a specified age interval [*t*_1_, *t* _2_] by computing the normalized integral: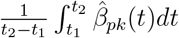 or 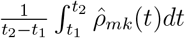 *dt*, where *t*_1_ and *t*_2_ denote the lower and upper bounds of the age interval, respectively. Lastly, to assess the contribution of informative genes to subtype differentiation, we computed odds ratios (ORs) for subtypes *Late-Onset Decline, Early Vulnerability* and *Rapid Decline* relative to *Resilient*. For a given signal gene *g*_*iq*_, the OR for subtype k (*k* = 2, 3, 4) relative to the *Resilient* subtype is calculated as 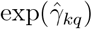.

#### Multimodal profiling reveals clinico-pathological and molecular divergence across TPClust subtypes

To evaluate differences in clinical, neuropathological, molecular, proteomic, and neuroimaging traits across TPClust-defined subtypes, we used generalized linear models (GLMs) adjusting for age and sex. Each variable was modeled as the response with subtype indicators, age, and sex as covariates, using the glm function in the R package stats (version 4.3.2). Pairwise subtype comparisons were conducted with the glht function in the R package multcomp (version 1.4.25), using custom contrasts specified to capture all relevant pairwise differences. Violin plots were generated to visualize subtype differences, accompanied by Wilcoxon and Kruskal–Wallis tests results using the stat_compare_means function from the ggplot2 package (version 3.5.1) in R.

#### T-distributed stochastic neighbor embedding (t-SNE) analysis for identified subtypes

To visualize subtype separation based on clinical and clinico-pathological profiles, we integrated AD-related traits with additional variables, including mean age, mean global cognition score, and diabetes, hypertension, and frailty. All continuous variables were normalized, and t-SNE algorithm was applied to reduce the data to two dimensions using the Rtsne function from the Rtsne package (version 0.17) in R.

#### Transcriptome-wide analyses

To identify distinct molecular pathways characterizing each TP-Clust subtype, we conducted transcriptome-wide differential gene expression (DGE) and GO BP enrichment analyses. DGE was conducted between subtypes, and enrichment analysis was performed using the clusterProfiler package (Yu et al., 2012), with false discovery rate (FDR) correction applied (FDR < 0.05).

#### Survival analysis

To evaluate subtype-specific differences in progression to clinical dementia, we conducted dementia-free survival analyses across TPClust-identified subtypes using an extended accelerated failure time (AFT) model (Jackson, 2016), accounting for left truncation since all ROSMAP participants were non-demented at baseline (Lee et al., 2023; Wilson et al., 2021). A generalized gamma distribution was specified for the survival time of the *Resilient* subtype, used as the reference group, to allow for subtype-specific time-varying hazard ratios. The analysis was implemented using the flexsurvreg function from the flexsurv R package (version 2.3.2).

To test for overall survival differences across subtypes, we used a Fleming–Harrington permutation test based on a weighted log-rank statistic. This approach offers a more robust and sensitive comparison of survival distributions under right-censoring compared to standard log-rank test. P-values were derived from 1,000 random permutations of subtype labels. Robustness was confirmed across a range of tuning parameters, all yielding p-values less than 0.001.

#### Benchmarking against unimodal subtyping methods

To compare the performance of TP-Clust’s integrative modeling with unimodal subtyping approaches, we evaluated its ability to resolve clinically and biologically meaningful heterogeneity. For the clinical-only subtyping, TPClust was applied using longitudinal global cognition data only, excluding omics features and associated feature selection regularizations, while retaining temporal smoothness regularization and other modeling components. The likelihood and EM algorithm were modified accordingly. Cross-validation was used to determine the optimal number of subtypes, yielding two: *Resilient clinic* and *Rapid Decline clinic*.

For the omics-only approach, we performed principal component analysis (PCA) on DLPFC gene expression data, retaining 308 principal components explaining 90% of the variance. K-means clustering was then applied, with the optimal number of subtypes determined using both silhouette score and elbow method, both indicating two subtypes. K-means algorithm was performed using the *kmeans* function from the base R package *stats* (version 4.3.2). To compare subtype assignments across methods, we used flow diagrams to visualize overlap between TPClust, clinical-only, and omics-only subtypes. These diagrams were generated using the plot_ly function from R package plotly (version 4.10.4).

## Author contributions

B.H. developed algorithms, performed data analyses, generated figures, interpreted results, and wrote the manuscript. B.N.V. interpreted ROSMAP results and reviewed the manuscript. P.L.D. provided ROSMAP data, contributed to result interpretation, and reviewed the manuscript. D.A.B. provided ROSMAP data and reviewed the manuscript. Y.W. conceived the study, developed algorithms, interpreted results, secured funding, and reviewed the manuscript. A.J.L. conceived the study, developed algorithms, performed data analyses, generated figures, interpreted results, secured funding, and wrote the manuscript.

## Data availability

The datasets used in this study are accessible through the Rush Alzheimer’s Disease Center (RADC) Research Resource Sharing Hub (https://www.radc.rush.edu), funded by the National Institute on Aging (NIA). This platform supports research on the causes, treatment, and prevention of AD and other chronic conditions of aging. Data access is available for research purposes in accordance with the data and resource request and sharing policy (https://www.radc.rush.edu/requests.htm).

## Code availability

The software developed in this work is publicly available at https://github.com/BoyiHu673/TPClust.

## Acknowledgements

This work was supported by the National Institutes of Health under grant nos. K01AG084849 (A.J.L.), P30AG06646 (A.J.L.), U19AG078109 (A.J.L.), R01NS073671 (Y.W.), NS073671 (Y.W.), and MH123487 (Y.W.), the National Alzheimer’s Coordinating Center–Alzheimer’s Association New Investigator Award (A.J.L.), and the Carol and Gene Ludwig Pilot Grant in Neurodegeneration (A.J.L.). ROSMAP is supported by P30AG10161, P30AG72975, R01AG17917, R01AG015819, U01AG072572, and U01AG046152. We acknowledge the participants of the Religious Orders Study and the Rush Memory and Aging Project, as well as the investigators and staff who collected and curated the data.

## Supplementary Tables

Table S1. Mean cognitive trajectories across TPClust-identified subtypes, adjusted for covariate prevalence

Table S2. Age-varying effects of covariates on cognition estimated by TPClust across TPClust-identified subtypes

Table S3. Dementia-free survival probabilities across TPClust-identified subtypes

Table S4. Cumulative risk of dementia across TPClust-identified subtypes

Table S5. Association analysis between the TPClust-identified subtypes and clinico-pathological traits

Table S6. Association analysis between TPClust-identified subtypes and clinical traits (MMSE, daily activity)

Table S7. Association analysis between TPClust-identified subtypes and brain infarcts

Table S8. Association analysis between TPClust-identified subtypes and cognitive decline after covariate adjustment

Table S9. Association analysis between TPClust-identified subtypes and molecular and cellular biomarkers

Table S10. Signal genes identified by TPClust and parameter estimates for selected genes

Table S11. Differential gene expression analysis comparing Rapid Decline and Resilient subtypes

Table S12. Differential gene expression analysis comparing Early Vulnerability and Resilient subtypes

Table S13. Differential gene expression analysis comparing Rapid Decline and Late-Onset Decline subtypes

Table S14. Differential gene expression analysis comparing Rapid Decline and Early Vulnerability subtypes

Table S15. GO biological process enrichment analysis comparing Rapid Decline and Resilient (FDR < 0.05)

Table S16. GO biological process enrichment analysis comparing Early Vulnerability and Resilient (FDR < 0.05)

Table S17. GO biological process enrichment analysis comparing Rapid Decline and Late-Onset Decline (FDR < 0.05)

Table S18. Association analysis between TPClust-identified subtypes and proteomic features

Table S19. Association analysis between TPClust-identified subtypes and MRI neuroimaging features

Table S20. Association analysis between subtypes identified using clinical data only and clinicopathological traits

Table S21. Association analysis between subtypes identified using omics data only and clinicopathological traits

Table S22. Differentially expressed genes between individuals with pathological AD and controls in ROSMAP (FDR < 0.05)

Table S23. Enriched Gene Ontology biological process pathways (FDR < 0.05)

## References

Amidei, C. B., A. Fayosse, J. Dumurgier, M. D. Machado-Fragua, A. G. Tabak, T. van Sloten, M. Kivimäki, A. Dugravot, S. Sabia, and A. Singh-Manoux (2021). Association between age at diabetes onset and subsequent risk of dementia. Jama 325(16), 1640–1649.

Arvanitakis, Z., S. E. Leurgans, L. L. Barnes, D. A. Bennett, and J. A. Schneider (2011). Microinfarct pathology, dementia, and cognitive systems. Stroke 42(3), 722–727.

Bellou, E., E. Baker, G. Leonenko, M. Bracher-Smith, P. Daunt, G. Menzies, J. Williams, V. Escott-Price, A. D. N. Initiative, et al. (2020). Age-dependent effect of apoe and polygenic component on alzheimer’s disease. Neurobiology of aging 93, 69–77.

Bennett, D. A., A. S. Buchman, P. A. Boyle, L. L. Barnes, R. S. Wilson, and J. A. Schneider (2018). Religious orders study and rush memory and aging project. Journal of Alzheimer’s disease 64(s1), S161–S189.

Buchman, A. S., P. A. Boyle, R. S. Wilson, Y. Tang, and D. A. Bennett (2007). Frailty is associated with incident alzheimer’s disease and cognitive decline in the elderly. Psychosomatic medicine 69(5), 483–489.

De Boor, C. (1986). B (asic)-spline basics. University of Wisconsin-Madison. Mathematics Research Center.

De Jager, P. L., Y. Ma, C. McCabe, J. Xu, B. N. Vardarajan, D. Felsky, H.-U. Klein, C. C. White, M. A. Peters, B. Lodgson, et al. (2018). A multi-omic atlas of the human frontal cortex for aging and alzheimer’s disease research. Scientific data 5(1), 1–13.

Felsky, D., I. Santa-Maria, M. I. Cosacak, L. French, J. A. Schneider, D. A. Bennett, P. L. De Jager, C. Kizil, and G. Tosto (2023). The caribbean-hispanic alzheimer’s disease brain transcriptome reveals ancestry-specific disease mechanisms. Neurobiology of disease 176, 105938.

Fleischman, D. A., K. Arfanakis, S. E. Leurgans, A. M. Evia, M. Lamar, A. Kapasi, S. D. Han, V. N. Poole, M. Wagner, D. A. Bennett, et al. (2025). Arts is associated with vascular risk factors, mci, dementia, and stroke. Alzheimer’s & Dementia 21(7), e70430.

Forgy, E. W. (1965). Cluster analysis of multivariate data: Efficiency versus interpretability of classifications. Biometrics 21(3), 768–769.

Grant, M., S. Boyd, and Y. Ye (2009). Cvx users’ guide. online: http://www.stanford.edu/boyd/software.html.

Gueniot, F., S. Rubin, P. Bougaran, A. Abelanet, J. L. Morel, B. Bontempi, C. Proust, P. Dufourcq, T. Couffinhal, and C. Duplaa (2022). Targeting pdzrn3 maintains adult blood-brain barrier and central nervous system homeostasis. Journal of Cerebral Blood Flow & Metabolism 42(4), 613–629.

Guo, J., M. Wall, and Y. Amemiya (2006). Latent class regression on latent factors. Biostatistics 7(1), 145–163.

Han, K., B. Kim, S. H. Lee, and M. K. Kim (2023). A nationwide cohort study on diabetes severity and risk of parkinson disease. npj Parkinson’s Disease 9(1), 11.

Heywood, A., J. Stocks, J. A. Schneider, K. Arfanakis, D. A. Bennett, M. F. Beg, and L. Wang (2022). The unique effect of tdp-43 on hippocampal subfield morphometry and cognition. NeuroImage: Clinical 35, 103125.

House, M. J., T. S. Pierre, J. Foster, R. Martins, and R. Clarnette (2006). Quantitative mr imaging r2 relaxometry in elderly participants reporting memory loss. American Journal of Neuroradiology 27(2), 430–439.

Jackson, C. H. (2016). flexsurv: a platform for parametric survival modeling in r. Journal of statistical software 70.

James, B. D., P. A. Boyle, D. A. Bennett, and A. S. Buchman (2012). Total daily activity measured with actigraphy and motor function in community-dwelling older persons with and without dementia. Alzheimer Disease & Associated Disorders 26(3), 238–245.

Lee, A. J., Y. Ma, L. Yu, R. J. Dawe, C. McCabe, K. Arfanakis, R. Mayeux, D. A. Bennett, H.-U. Klein, and P. L. De Jager (2023). Multi-region brain transcriptomes uncover two subtypes of aging individuals with differences in alzheimer risk and the impact of apoeε4. bioRxiv, 2023–01.

Lee, A. J., D. Sanchez, D. Reyes-Dumeyer, A. M. Brickman, R. A. Lantigua, B. N. Vardarajan, and R. Mayeux (2023). Reliability and validity of self-reported vascular risk factors: Hypertension, diabetes, and heart disease, in a multi-ethnic community based study of aging and dementia. Journal of Alzheimer’s Disease Preprint, 1–11.

Li, Q. S. and L. De Muynck (2021). Differentially expressed genes in alzheimer’s disease highlighting the roles of microglia genes including olr1 and astrocyte gene cdk2ap1. Brain, Behavior, & Immunity-Health 13, 100227.

Li, Y., P. Liu, W. Wang, W. Zong, Y. Fang, Z. Ren, L. Tang, J. C. Celedón, S. Oesterreich, and G. C. Tseng (2024). Outcome-guided disease subtyping by generative model and weighted joint likelihood in transcriptomic applications. The Annals of Applied Statistics 18(3), 1947–1964.

Ma, Y., L. Yu, M. Olah, R. Smith, S. R. Oatman, M. Allen, E. Pishva, B. Zhang, V. Menon, N. Ertekin-Taner, et al. (2022). Epigenomic features related to microglia are associated with attenuated effect of apoe ε4 on alzheimer’s disease risk in humans. Alzheimer’s & Dementia 18(4), 688–699.

Meier, L., S. Van De Geer, and P. Bühlmann (2008). The group lasso for logistic regression. Journal of the Royal Statistical Society Series B: Statistical Methodology 70(1), 53–71.

Meinshausen, N. (2007). Relaxed lasso. Computational Statistics & Data Analysis 52(1), 374–393.

Meng, L., D. Avram, G. Tseng, and Z. Huo (2022). Outcome-guided sparse k-means for disease subtype discovery via integrating phenotypic data with high-dimensional transcriptomic data. Journal of the Royal Statistical Society Series C: Applied Statistics 71(2), 352–375.

Meng, X.-L. and D. Van Dyk (1997). The em algorithm—an old folk-song sung to a fast new tune. Journal of the Royal Statistical Society Series B: Statistical Methodology 59(3), 511–567.

Miao, J., Y. Zhang, C. Su, Q. Zheng, and J. Guo (2025). Insulin-like growth factor signaling in alzheimer’s disease: Pathophysiology and therapeutic strategies. Molecular Neurobiology 62(3), 3195–3225.

Milind, N., C. Preuss, A. Haber, G. Ananda, S. Mukherjee, C. John, S. Shapley, B. A. Logsdon, P. K. Crane, and G. W. Carter (2020). Transcriptomic stratification of late-onset alzheimer’s cases reveals novel genetic modifiers of disease pathology. PLoS genetics 16(6), e1008775.

Neff, R. A., M. Wang, S. Vatansever, L. Guo, C. Ming, Q. Wang, E. Wang, E. Horgusluoglu-Moloch, W.-m. Song, A. Li, et al. (2021). Molecular subtyping of alzheimer’s disease using rna sequencing data reveals novel mechanisms and targets. Science advances 7(2), eabb5398.

Olah, M., E. Patrick, A.-C. Villani, J. Xu, C. C. White, K. J. Ryan, P. Piehowski, A. Kapasi, P. Nejad, M. Cimpean, et al. (2018). A transcriptomic atlas of aged human microglia. Nature communications 9(1), 539.

Proust-Lima, C., M. Séne, J. M. Taylor, and H. Jacqmin-Gadda (2014). Joint latent class models for longitudinal and time-to-event data: a review. Statistical methods in medical research 23(1), 74–90.

Ramos-Miguel, A., A. A. Jones, V. A. Petyuk, V. E. Barakauskas, A. M. Barr, S. E. Leurgans, P. L. De Jager, K. B. Casaletto, J. A. Schneider, D. A. Bennett, et al. (2021). Proteomic identification of select protein variants of the snare interactome associated with cognitive reserve in a large community sample. Acta Neuropathologica 141, 755–770.

Ramsay, J. and B. Silverman (2005). Functional Data Analysis (2nd ed.). Springer Series in Statistics. New York: Springer.

Sahelijo, N., P. Rajagopalan, L. Qian, R. Rahman, D. Priyadarshi, D. Goldstein, S. I. Thomopoulos, D. A. Bennett, L. A. Farrer, T. D. Stein, et al. (2024). Brain cell-based genetic subtyping and drug repositioning for alzheimer disease. medRxiv, 2024–06.

Simon, N., J. Friedman, T. Hastie, and R. Tibshirani (2013). A sparse-group lasso. Journal of computational and graphical statistics 22(2), 231–245.

Sullivan, S. E., M. Liao, R. V. Smith, C. White, V. N. Lagomarsino, J. Xu, M. Taga, D. A. Bennett, P. L. De Jager, and T. L. Young-Pearse (2019). Candidate-based screening via gene modulation in human neurons and astrocytes implicates fermt2 in a β and tau proteostasis. Human molecular genetics 28(5), 718–735.

Sun, J., J. D. Herazo-Maya, P. L. Molyneaux, T. M. Maher, N. Kaminski, and H. Zhao (2019). Regularized latent class model for joint analysis of high-dimensional longitudinal biomarkers and a time-to-event outcome. Biometrics 75(1), 69–77.

Tang, M.-X., Y. Stern, K. Marder, K. Bell, B. Gurland, R. Lantigua, H. Andrews, L. Feng, B. Tycko, and R. Mayeux (1998). The apoe-epsilon4 allele and the risk of alzheimer disease among african americans, whites, and hispanics. Jama 279(10), 751–755.

Tibshirani, R. (1996). Regression shrinkage and selection via the lasso. Journal of the Royal Statistical Society Series B: Statistical Methodology 58(1), 267–288.

Trzepacz, P. T., H. Hochstetler, S. Wang, B. Walker, A. J. Saykin, and A. D. N. Initiative (2015). Relationship between the montreal cognitive assessment and mini-mental state examination for assessment of mild cognitive impairment in older adults. BMC geriatrics 15(1), 107.

Van Der Vaart, A. W. and J. A. Wellner (1996). Weak convergence. In Weak convergence and empirical processes: with applications to statistics, pp. 16–28. Springer.

Wilson, R. S., P. A. Boyle, L. Yu, L. L. Barnes, J. Sytsma, A. S. Buchman, D. A. Bennett, and J. A. Schneider (2015). Temporal course and pathologic basis of unawareness of memory loss in dementia. Neurology 85(11), 984–991.

Wilson, R. S., T. Wang, L. Yu, F. Grodstein, D. A. Bennett, and P. A. Boyle (2021). Cognitive activity and onset age of incident alzheimer disease dementia. Neurology 97(9), e922–e929.

Yu, G., L.-G. Wang, Y. Han, and Q.-Y. He (2012). clusterprofiler: an r package for comparing biological themes among gene clusters. Omics: a journal of integrative biology 16(5), 284–287.

Yu, L., P. Boyle, R. S. Wilson, E. Segawa, S. Leurgans, P. L. De Jager, and D. A. Bennett (2012). A random change point model for cognitive decline in alzheimer’s disease and mild cognitive impairment. Neuroepidemiology 39(2), 73–83.

Yu, L., P. A. Boyle, E. Segawa, S. Leurgans, J. A. Schneider, R. S. Wilson, and D. A. Bennett (2015). Residual decline in cognition after adjustment for common neuropathologic conditions. Neuropsychology 29(3), 335.

Yu, L., V. A. Petyuk, C. Gaiteri, S. Mostafavi, T. Young-Pearse, R. C. Shah, A. S. Buchman, J. A. Schneider, P. D. Piehowski, R. L. Sontag, et al. (2018). Targeted brain proteomics uncover multiple pathways to alzheimer’s dementia. Annals of neurology 84(1), 78–88.

Yu, L., V. A. Petyuk, K. d. P. Lopes, S. Tasaki, V. Menon, Y. Wang, J. A. Schneider, P. L. De Jager, and D. A. Bennett (2023). Associations of vgf with neuropathologies and cognitive health in older adults. Annals of neurology 94(2), 232–244.

Zou, H. and T. Hastie (2005). Regularization and variable selection via the elastic net. Journal of the Royal Statistical Society Series B: Statistical Methodology 67(2), 301–320.

