## Supplementary Information for "TPClust: Temporal Profile-Guided Subtyping Using High-Dimensional Omics Data"

#### Contents

**Supplementary Note 1. Computational Cost of TPClust in Real Application** **2**

**Supplementary Note 2. Multimodal profiling reveals clinico-pathological and molecular divergence across TPClust subtypes** **2**

**Supplementary Note 3. Integrative modeling in TPClust outperforms unimodal subtyping approaches** **3**

**Supplementary Note 4. Algorithm for Model Fitting.** **4**

**Supplementary Note 5. Simulation Study** **5**

**Supplementary Tables.** **6**

### Supplementary Note 1. Computational Cost of TPClust in Real Application

To assess the computational cost of TPClust, we evaluated its runtime on subsets of the ROSMAP dataset with sample sizes of 200 and 500. Each subset was randomly drawn from the full dataset and included 2,015 transcriptomic features and six covariates. Computations were performed on high-performance computing clusters equipped with Intel(R) Xeon(R) CPU E5-2620 v4 @ 2.10GHz processors. For subsets with 200 samples, the runtime of TPClust under fixed tuning parameter settings ranged from approximately 20 minutes to 1 hour, depending on the initialization of the EM algorithm and the choice of tuning parameters. For subsets with sample size 500, the runtime ranged from approximately 100 minutes to about 6 hours. When applied to the full ROSMAP dataset, model fitting completed in 12 -36 hours using 96 CPU cores. The number of cores utilized depended on the number of clusters considered and the number of candidate values specified for each tuning parameter. In addition, we evaluated the runtime per EM iteration on a MacBook Pro equipped with Apple M3 chips. For the full ROSMAP dataset, the duration of a single EM iteration ranged from 5.77 to 37.15 minutes, depending on the number of clusters, initialization strategy, and tuning parameter choices, with an average runtime of approximately 16 minutes.

### Supplementary Note 2. Multimodal profiling reveals clinico-pathological and molecular divergence across TPClust subtypes

For proteomics analysis, in addition to the major findings in the main text, the *Late-onset Decline* subtype also showed intermediate profiles with elevated synaptic proteins (*SNAP25*, *STX1A*,

*SYT12*, and *VGF*) and reduced increased $\beta$-amyloid (*bA*), stress-related protein (*IGFBP5*) and glial and inflammatory markers (*HSPB2*). For neuroimaging features, the *Rapid Decline* subtype exhibited significantly reduced cortical thickness in 24 brain regions, including AD-vulnerable areas such as the entorhinal cortex, fusiform gyrus, inferior parietal lobule, inferior temporal gyrus, and parahippocampal gyrus. Gray matter volumes were also significantly lower in 27 regions spanning limbic and neocortical areas, including the hippocampus, amygdala, and caudal middle frontal gyrus.

### Supplementary Note 3. Integrative modeling in TPClust outperforms unimodal subtyping approaches

To extend the transcriptomics-only comparison, we evaluated two clusters identified by integrating gene expression across three brain regions in ROSMAP using sparse canonical correlation analysis followed by K-means clustering (Lee et al., 2023). The resulting meta-clusters (MC1 and MC2) closely resembled the PCA-based subtypes but showed limited alignment with TPClust subtypes (Fig. 5c). MC2 included both *Resilient* (62.1%) and *Rapid Decline* (13.9%) individuals, while MC1 comprised 21.6% *Rapid Decline* and 48.6% *Resilient* cases (Fig. 5d), obscuring the clinico-pathological distinctions captured by TPClust.

### Supplementary Note 4. Algorithm for Model Fitting

For each $k = 1, \ldots K$, define

$$\boldsymbol{Y} = \begin{pmatrix} y_{11} \\ y_{12} \\ \ldots \\ y_{1T_1} \\ y_{21} \\ \ldots \\ y_{nT_n} \end{pmatrix}, \quad \boldsymbol{\alpha} = \begin{pmatrix} \alpha_{1k} \\ \alpha_{1k} \\ \ldots \\ \alpha_{1k} \\ \alpha_{2k} \\ \ldots \\ \alpha_{nk} \end{pmatrix}, \quad \boldsymbol{W}_k = \begin{pmatrix} \sqrt{w_{1k}^{(s)}} & 0 & \ldots & \ldots \\ 0 & \sqrt{w_{1k}^{(s)}} & \ldots & \\ \ldots & \ldots & \ldots & \ldots \\ \ldots & \ldots & 0 & \sqrt{w_{nk}^{(s)}} \end{pmatrix},$$

$$\boldsymbol{X} = \begin{pmatrix} x_{10}\boldsymbol{b}(t_{11})^\tau & \ldots & x_{1p}\boldsymbol{b}(t_{11})^\tau & v_{11}(t_{11})\boldsymbol{b}(t_{11})^\tau & \ldots & v_{1M}(t_{11})\boldsymbol{b}(t_{11})^\tau \\ x_{10}\boldsymbol{b}(t_{12})^\tau & \ldots & x_{1p}\boldsymbol{b}(t_{12})^\tau & v_{11}(t_{12})\boldsymbol{b}(t_{12})^\tau & \ldots & v_{1M}(t_{12})\boldsymbol{b}(t_{12})^\tau \\ \ldots & \ldots & \ldots & \ldots & \ldots & \ldots \\ x_{10}\boldsymbol{b}(t_{1T_1})^\tau & \ldots & x_{1p}\boldsymbol{b}(t_{1T_1})^\tau & v_{11}(t_{1T_1})\boldsymbol{b}(t_{1T_1})^\tau & \ldots & v_{1M}(t_{1T_1})\boldsymbol{b}(t_{1T_1})^\tau \\ x_{10}\boldsymbol{b}(t_{21})^\tau & \ldots & x_{1p}\boldsymbol{b}(t_{21})^\tau & v_{11}(t_{21})\boldsymbol{b}(t_{21})^\tau & \ldots & v_{1M}(t_{21})\boldsymbol{b}(t_{21})^\tau \\ \ldots & \ldots & \ldots & \ldots & \ldots & \ldots \\ x_{n0}\boldsymbol{b}(t_{nT_n})^\tau & \ldots & x_{1p}\boldsymbol{b}(t_{nT_n})^\tau & v_{11}(t_{nT_n})\boldsymbol{b}(t_{nT_n})^\tau & \ldots & v_{nM}(t_{nT_n})\boldsymbol{b}(t_{nT_n})^\tau \end{pmatrix},$$

and

$$\boldsymbol{U} = \begin{pmatrix} \nu_1\boldsymbol{Q} & \boldsymbol{0}^\tau & \ldots & \boldsymbol{0}^\tau \\ \boldsymbol{0} & \nu_2\boldsymbol{Q} & \ldots & \boldsymbol{0}^\tau \\ \ldots & \ldots & \ldots & \ldots \\ \boldsymbol{0} & \ldots & \ldots & \kappa_M\boldsymbol{Q} \end{pmatrix}.$$

### Supplementary Note 5. Simulation Study.

Let $\boldsymbol{b}(t)$ denote the B-splines defined over $[0,1]$ with 8 equally spaced knots and order equal to 4. The total number of B-splines is 10. Let $\boldsymbol{c}_{01}, \boldsymbol{c}_{11}, \ldots, \boldsymbol{c}_{23}$ denote the vectors of coefficients with same length as $\boldsymbol{b}(t)$ ranging from $-1.5$ to $2.5$. The time-varying coefficients $\beta_{jk}(t)$, $j = 0,1,2$ and $k = 1,2,3$, are defined as

$$\beta_{jk}(t) = \boldsymbol{c}_{jk}^{\tau}\boldsymbol{b}(t), \quad j = 0,1,2, \quad k = 1,2,3.$$

The coefficients $\rho_{1,k}$ are defined as

$$\rho_{11}(t) = -t, \quad \rho_{12}(t) = t, \quad \rho_{13}(t) = 2t^2.$$

The random effects $\alpha_{ik}$ are generated from two different normal distributions:

$$\alpha_{i1} \overset{\text{IID}}{\sim} \text{N}(0, 0.5^2), \quad \alpha_{i2} \overset{\text{IID}}{\sim} \text{N}(0, 1), \quad \alpha_{i3} \overset{\text{IID}}{\sim} \text{N}(0, 0.5^2).$$

The random errors are generated from three different normal distributions:

$$\epsilon_{ij1} \overset{\text{IID}}{\sim} N(0, 0.5^2), \quad \epsilon_{ij2} \overset{\text{IID}}{\sim} N(0, 0.3^2), \quad \epsilon_{ij1} \overset{\text{IID}}{\sim} N(0, 0.2^2).$$

To mimic the phenotype data, we generated two binary covariates for each individual:

$$X_{i1} \overset{\text{IID}}{\sim} \text{Bernoulli}(0.4), \quad X_{i2} \overset{\text{IID}}{\sim} \text{Bernoulli}(0.6).$$

To mimic clinical data with persistence over time, we generated a time-varying binary variable $V_{i1}(t)$ from a Bernoulli distribution at each time point $t$:

$$V_{i1}(t) \sim \text{Bernoulli}(0.3).$$

Once $V_{i1}(t) = 1$ at any time point $t$, all subsequent observations for that individual were set to 1, i.e., $V_{i1}(t') = 1$ for all $t' > t$. For the generation of omics data, the feature vector $\boldsymbol{g}_i$ was generated from independent and identically distributed uniform distributions:

$$\boldsymbol{g}_i \overset{\text{i.i.d.}}{\sim} \text{Unif}(-0.5, 0.5).$$

For the coefficient vector $\boldsymbol{\gamma}_2$, the non-zero entries were set to values ranging from 1 to 3. For $\boldsymbol{\gamma}_3$, the non-zero entries ranged from $-5$ to $-2$.

#### Supplementary Tables.

Supplementary tables S1-S23 are saved in the Excel document named "Supplementary Tables".

#### References

Lee, A. J., Y. Ma, L. Yu, R. J. Dawe, C. McCabe, K. Arfanakis, R. Mayeux, D. A. Bennett, H.-U. Klein, and P. L. De Jager (2023). Multi-region brain transcriptomes uncover two subtypes of aging individuals with differences in alzheimer risk and the impact of apoe$\varepsilon$4. *bioRxiv*, 2023–01.
